# AnnoSpat annotates cell types and quantifies cellular arrangements from spatial proteomics

**DOI:** 10.1101/2023.01.15.524135

**Authors:** Aanchal Mongia, Diane C. Saunders, Yue J. Wang, Marcela Brissova, Alvin C. Powers, Klaus H. Kaestner, Golnaz Vahedi, Ali Naji, Gregory W. Schwartz, Robert B. Faryabi

## Abstract

Cellular composition and anatomical organization influence normal and aberrant organ functions. Emerging spatial single-cell proteomic assays such as Image Mass Cytometry (IMC) and Co-Detection by Indexing (CODEX) have facilitated the study of cellular composition and organization by enabling high-throughput measurement of cells and their localization directly in intact tissues. However, annotation of cell types and quantification of their relative localization in tissues remain challenging. To address these unmet needs, we developed AnnoSpat (Annotator and Spatial Pattern Finder) that uses neural network and point process algorithms to automatically identify cell types and quantify cell-cell proximity relationships. Our study of data from IMC and CODEX show the superior performance of AnnoSpat in rapid and accurate annotation of cell types compared to alternative approaches. Moreover, the application of AnnoSpat to type 1 diabetic, non-diabetic autoantibody-positive, and non-diabetic organ donor cohorts recapitulated known islet pathobiology and showed differential dynamics of pancreatic polypeptide (PP) cell abundance and CD8^+^ T cells infiltration in islets during type 1 diabetes progression.

## Introduction

Tissues consist of diverse cell types whose functions are influenced by communication and interaction with surrounding cells. In addition to cell intrinsic aberrations, dysfunction in the cellular microenvironment impacts organ function and contributes to pathology of complex diseases, such as type 1 diabetes. The emergence of spatially resolved single-cell proteomic assays such as Image Mass Cytometry (IMC) and Co-Detection by Indexing (CODEX) has allowed high-throughput measurement of cellular composition and localization within intact tissues and advanced understanding of intricate cell-cell interactions. However, the unique characteristics of spatial proteomic assays, coupled with their ability to measure millions of cells, have created a need for efficient and automated computational tools that enable identification of cell-types and quantification of their spatial colocalization. To address this unmet need, we introduce AnnoSpat (Annotator and Spatial Pattern Finder) for rapid, scalable, and automated annotation of cell types and quantification of their spatial relationships.

Despite the paucity of algorithms for cell-type annotation from IMC and CODEX data, several algorithms have been proposed to predict cell types from single-cell RNA sequencing (scRNA-seq) data^1^. Many of these methods, such as scmap and Garnett, use clustering to group together transcriptionally similar cells and then map each cluster to reference cell types from *a priori* annotated datasets using representative cells from each group^2,3^. These methods rely on accurate clustering and reference data annotation, which was previously characterized based on manual assessment of differential expression of selected marker genes. Another category of scRNA-seq cell-type annotators use supervised machine learning models such as support vector machines^4^, neural networks^5^, and random forests^6,7^. Similarity-based methods, such as TooManyPeaks^8^, are the third category of methods that annotate cell types based on bulk measurement of purified reference cell populations. Training of supervised machine learning- and similarity-based methods require large sets of purified or expert-annotated cell populations, which are respectively lacking for *in situ* proteomic assays such as IMC and CODEX. Unique characteristics of IMC and CODEX data further limit the use of existing cell-type annotation methods developed for scRNA-seq. While scRNA-seq experiments provide expression of thousands of genes for cell type prediction, IMC and CODEX measure the expression of tens of proteins. Furthermore, IMC and CODEX readouts are continuous intensities that cannot be readily inputted to most scRNA-seq cell-type annotators, such as Garnett, that only accept scRNA-seq count readouts. To address such limitations, Astir was recently proposed specifically for cell-type annotation from IMC data^9^. This method uses deep recognition neural networks for inference of cell types based on known marker proteins. Benchmarking of Astir suggests that supervised- and marker-based cell-type annotation methods tend to outperform other approaches^9^. Guided by this observation, we developed AnnoSpat by combining semi-supervised and supervised learning methods for cell-type annotation of IMC and CODEX data in the absence of manually labeled cells for training.

Cell-type annotation is an initial step in the analysis of most spatial proteomic data such as IMC and CODEX. To fully benefit from *in situ* single-cell assays and investigate tissue microenvironment, methods are needed to quantify the spatial organization of cells in regions of interest. The current methods either measure cell density across distances^7^, use Bayesian models estimating cell types across locations^10^, or use Ripley statistics^11^. To create a comprehensive tool capable of automating annotation of cell types and quantifying their spatial relationships, we also equipped AnnoSpat with new point process-based algorithms that relate not only the distribution of a single cell type in a region of interest as with Ripley’s K function statistics, but also examine the interaction of multiple cell types.

We assessed the accuracy and efficiency of AnnoSpat by benchmarking its ability to identify various cell types within pancreatic tissues. In addition to quantitative comparative benchmarking using IMC and CODEX data, we evaluated AnnoSpat performance based on expert annotated pancreata of type 1 diabetes (T1D) and non-diabetic donors. Given that pancreas is the site of T1D pathogenesis in which the host immune system mounts a response to insulin-secreting pancreatic beta cells, we further used AnnoSpat to study the microenvironment of pancreata from donors with autoantibodies towards pancreatic islet proteins in their blood but no clinical diagnosis of T1D (AAb^+^) to better understand T1D progression. Together, our comprehensive analysis of 1,170,000 cells from 143 slides of 19 Human Pancreas Analysis Program (HPAP) donors revealed the effectiveness of AnnoSpat in reliably identifying cell types and quantifying their spatial organization in complex tissues. AnnoSpat and its individual components are available through https://github.com/faryabiLab/AnnoSpat.

## Results

### AnnoSpat identifies cell types and quantifies their relative localization

To predict the identity of individual cells and quantify their localization within tissues, we developed AnnoSpat for automated analysis of spatially aware single-cell proteomic data (Figure 1). AnnoSpat provides an end-to-end solution for analysis of IMC and CODEX data (Figure 1A) by implementing two distinct but complementary functionalities: “Annotator” (Figure 1B) and “Spatial Pattern Finder” (Figure 1C).

**Figure 1:**
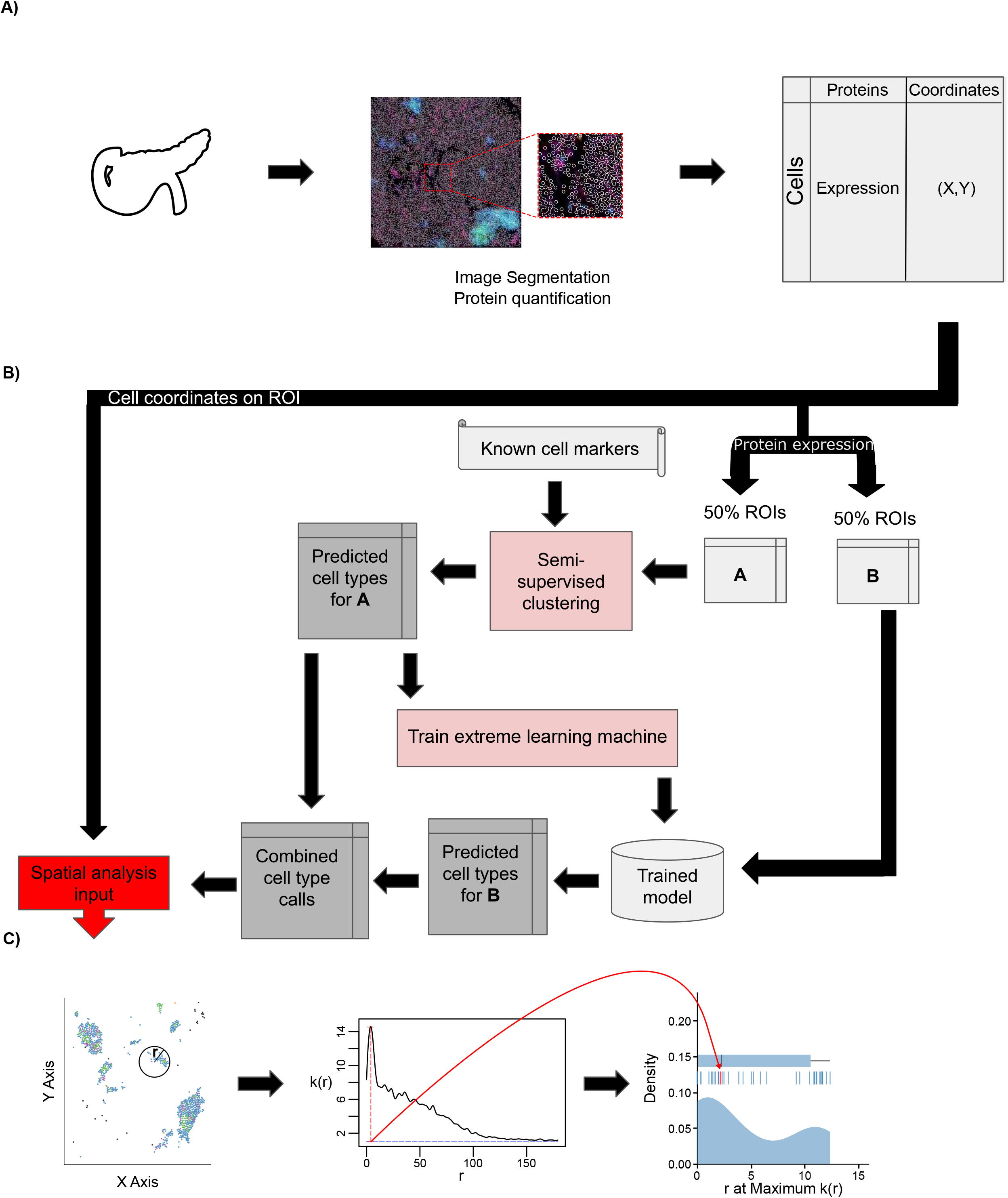
Overview of IMC or CODEX data analysis with AnnoSpat (Annotator and Spatial Pattern Finder). **(A)** From left to right: A tissue’s region of interest (ROI) (e.g. from the pancreas) is measured using a spatial single-cell proteomics assay such as IMC or CODEX, reporting position and protein expression levels of individual cells *in situ*. **(B)** To overcome lack of manually annotated training data, AnnoSpat’s Annotator module learns a cell-type predictor by first processing protein expression data with a semi-supervised clustering algorithm, which creates a training dataset from a subset of cells in the overall dataset (e.g. 50% in matrix A). Using this automatically generated training data, AnnoSpat then trains and applies an extreme learning machine classifier to label the remaining cells (e.g. 50% in matrix B). **(C)** AnnoSpat’s Spatial Pattern Finder component interprets cell locations as point processes to quantify relationships between cell types using distance-dependent (*r*) mark cross-correlation function (*k*(*r*)). Mark cross-correlation functions across ROIs are systematically summarized using different features of them such as the distance where the function is maximal.

To address the unmet need for annotating individual IMC- or CODEX-measured cells, the Annotator module of AnnoSpat learns a cell-type predictor from the matrix of raw protein expression levels and a list of *a priori* cell-type marker proteins. To overcome the lack of manually annotated training data, AnnoSpat implements a two-step learning process (Figure 1B). First, AnnoSpat deploys a constrained K-means semi-supervised clustering algorithm to create training data from a subset of cells in the dataset. Using this automatically generated training data, AnnoSpat then trains a classifier that will be used to predict the identity of additional cells. The number of clusters is set to the number of expected cell types within the tissue of interest along with an optional “Unknown” group that could account for cell types omitted from the marker protein list. To enhance the accuracy of K-means clustering, AnnoSpat initializes each cluster with cells that were annotated with high confidence based on distinct expression of marker proteins (Materials and Methods). This crucial step provides semi-supervision to the clustering algorithm, guiding AnnoSpat in grouping a subset of cells with similar protein expression levels into cell-type-labeled training cells. Taking this automatically generated training data, AnnoSpat then learns an extreme learning machine (ELM) classifier. ELM is a feed-forward neural network with non-iterative single step learning, which does not require tuning and backpropagation, and provides generalization performance and orders of magnitude faster learning compared to support vector machines and multi-layer perceptron^12^ (Materials and Methods). Together, the two-step learning algorithm equips AnnoSpat with an efficient and accurate cell annotation mechanism.

To enable the study of tissue microenvironment, we equipped AnnoSpat with the Spatial Pattern Finder module, which takes as input the Annotator-predicted cell types and their physical coordinates on the tissue region of interest (ROI) and quantifies cellular localization patterns (Figure 1C). The Spatial Pattern Finder algorithm applies point process theory to summarize cell relationships across a range of distances, from local neighborhoods to remote cells. Briefly, AnnoSpat compares cell pairs based on their cell type to any randomly chosen cells at a given distance apart. This process returns a mark cross-correlation function, a measure of cell-type aggregation at different distances (see Materials and Methods). The application of the mark cross-correlation function across ROIs allows for systematic quantification and comparison of inter-cell-type proximity in different conditions (Figure 1C). In addition to AnnoSpat software, we implemented Spatial Pattern Finder within the TooManyCells single-cell analysis suite^13^. This implementation includes the generation of interactive proximity plots that may be filtered by protein expression to fine-tune cell-type annotation. These interactive features also assist with exploration of spatial cell relationships. AnnoSpat’s Annotator and Spatial Pattern Finder functionalities together provide a solution for rapid and accurate annotation of millions of cells to study tissue microenvironment and cellular organization.

### AnnoSpat accurately identifies cell types in complex pancreatic tissues

To assess AnnoSpat’s Annotator performance, we first used IMC experiments measuring 33 proteins in pancreata from T1D and non-diabetic donors (Tables S1 and S2), and compared the ability of AnnoSpat, semi-supervised clustering (SSC), SCINA, AUCell, and Astir in identifying endocrine cell types. We considered these methods for comparative analysis since similar to AnnoSpat, they automate cell-type annotation and do not need training data. Astir uses a probabilistic Bayesian framework and is the only method developed for cell-type annotation from proteomics data^9^. SSC is a variant of AnnoSpat with its classifier replaced by centroids from the semi-supervised clustering step in Figure 1B. SCINA^14^ and AUCell^15^ use expectation-maximization and gene expression ranking for cell-type annotation from scRNA-seq count data, respectively. Default or suggested filters and parameters were used for all algorithms except AUCell, where size-factor normalization was disabled due to differences between the characteristics of discrete scRNA-seq count and continuous IMC data. Here, we used the canonical protein markers listed in Table S3 as an input to AnnoSpat and SCC.

To examine the extent of protein expression homogeneity in cell types predicted by these cell-type annotation methods, we compared their performances on 10 sets of 50, 000 randomly selected cells using the Silhouette Index (SI) and Davies Bouldin (DB) metrics. While SI assesses how a cell’s protein expression differs from other cells assigned the same type versus those assigned other types, DB reports the average similarity of each cell type with its most similar cell type, where similarity is defined as the ratio of intra-cell-type to inter-cell-type protein expression distances. More accurate cell-type annotation results in higher SI and lower DB. Based on these metrics, we observed differential performance of algorithms that depended on both cell type and disease status (Figures 2A, 2B, and 2C row 1; Tables S4, S5, and S6; and Supplemental Notes for AnnoSpat Benchmarking). For instance, AnnoSpat and SSC more accurately detected delta cells in the control samples compared to other algorithms. Notably, Astir, developed for cell-type annotation from IMC data, showed lower accuracy in identifying many cell types in both control and T1D samples.

**Figure 2:**
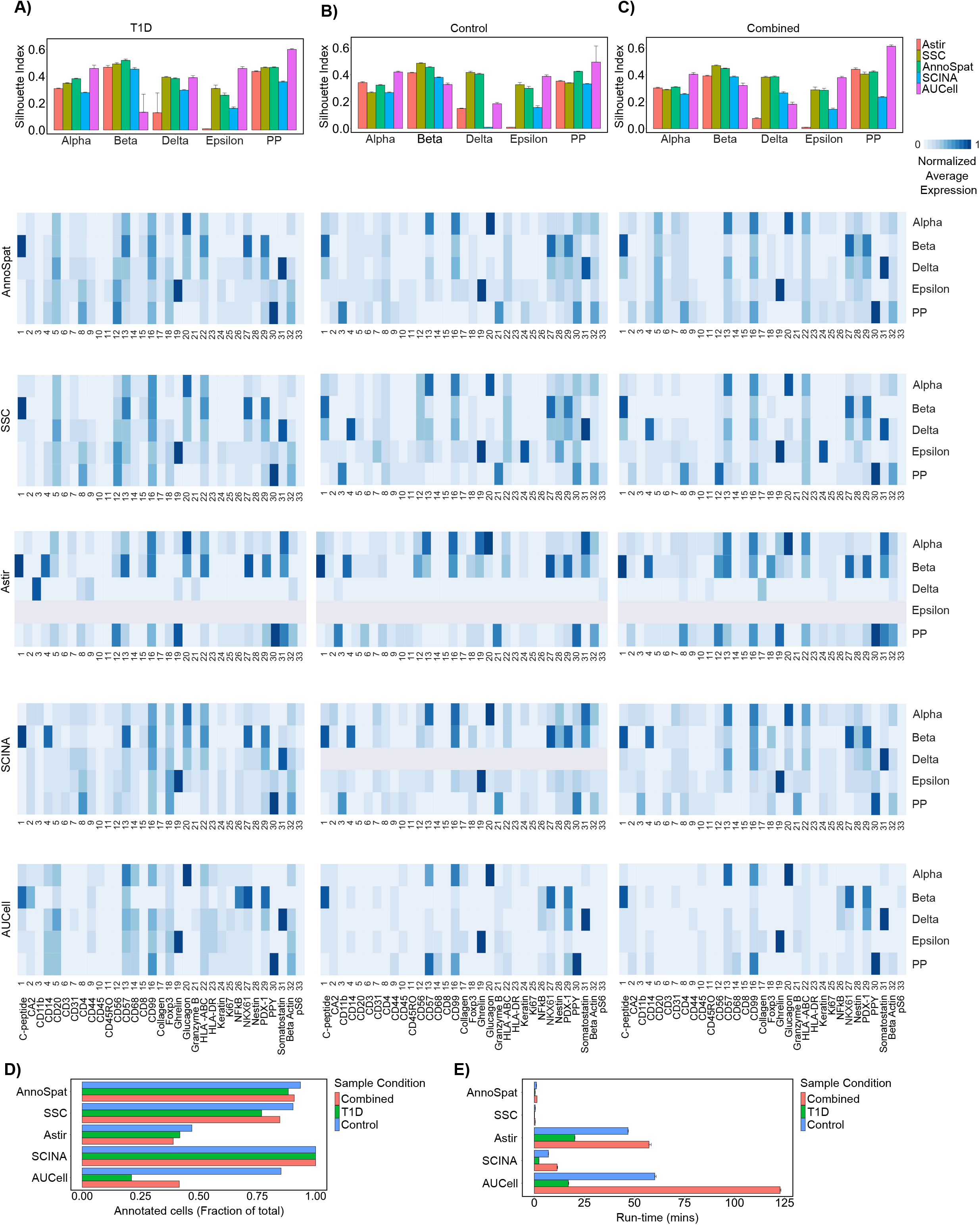
Comparative analysis of AnnoSpat cell-type annotation from IMC data. **(A)** From top to bottom: bar plots with error bars showing average and standard deviation Silhouette Index (SI), heatmaps showing marker proteins’ normalized average expressions for cells annotated as alpha, beta, delta, epsilon, and PP by AnnoSpat, semi-supervised clustering (SSC), Astir, SCINA, and AUCell from T1D pancreas IMC data (*n* = 374, 397 measured cells). **(B)** Similar to (A) from non-diabetic (control) pancreas IMC data (*n* = 795, 604 measured cells). **(C)** Similar to (A) from combined T1D and control pancreas IMC data (*n* = 1, 170, 001 measured cells). *m* = 10 sets of *n* = 50, 000 randomly selected cells are used for evaluation using SI in each bar plot in top panel of A-C. **(D)** Bar plots showing the fraction of *n* = 374, 397, *n* = 795, 604, and *n* = 1, 170, 001 IMC-measured cells form T1D, control, and combined T1D and control pancreata, respectively, annotated by AnnoSpat, SSC, Astir, SCINA, and AUCell. **(E)** Bar plots with error bars showing mean and standard deviation of run-time for the listed algorithms to annotate cells as in (D). Each algorithm was run three times on a machine with Ubuntu 20.04, 1.05TB Memory, Intel Xeon Gold CPU 6230R @ 2.1GHz, 2 physical processors 52 cores, and 104 threads.

To complement quantitative benchmarking and further evaluate the performance of cell-type annotation algorithms, we inspected protein expression profiles of cells labeled as alpha, beta, pancreatic polypeptide (PP), delta, and epsilon from IMC of T1D and non-diabetic donors. Compared to other cell types, these endocrine populations were particularly suitable for comparative analysis due to higher quality of their antibodies. We used a variant of term-frequency inverse document frequency (TF-IDF) normalization to reduce the effect of non-specific antibodies such as anti-CD99 and anti-beta actin on data visualization (Figure S1 and Materials and Methods). Inspection of endocrine canonical marker protein expression confirmed our quantitative bench-marking (Figures 2A, 2B, and 2C row 1) and showed the higher performance of AnnoSpat compared to other algorithms (Figures 2A, 2B, and 2C, rows 2 to 6; and Supplemental Notes for AnnoSpat Benchmarking). In addition to endocrine cells, AnnoSpat effectively detected other cell types that had high quality antibodies and are commonly present in the pancreatic tissue (Figures S1C, S2, and Supplemental Notes for AnnoSpat Benchmarking).

In addition to accuracy, we compared completeness and run-time of cell-type annotation. By using ELM, AnnoSpat annotated more than 90% of 1.1 million cells (Figure 2D and Table S7) in less than 2 minutes, a run-time only 3-times longer than SSC and notably faster than all other compared algorithms (Figure 2E and Table S8). Although SSC and AnnoSpat mostly exhibited comparable performance, close examination of data highlighted the additional benefit of AnnoSpat (Figures 2A, 2B, and 2C, rows 1 and 2, and S1). For instance, SSC-but not AnnoSpat-annotated delta cells expressed high levels of CD14, a protein expressed in macrophages and not delta cells (Figures 2B and 2C, rows 1 and 2). Notably, Astir failed to label nearly 50% of the cells (Figure 2D, Table S7) while took 40-times longer (Figure 2E, Table S8). Due to its bi-modal distribution model, SCINA assigned a label to almost all the cells in a reasonable time (Figures 2D and 2E, Tables S7 and S8) at the expense of diminished accuracy (Figures 2A, 2B, and 2C, rows 1, 2 and 5). Conversely, AUCell exhibited comparable performance to AnnoSpat (Figures 2A, 2B, and 2C, rows 1, 2 and 6), but it failed to annotate most cells included in the benchmarking analysis, potentially leading to information loss. Close examination of data revealed that AUCell more accurately labeled cell types with a larger number of marker proteins such as ductal cells (Figure S2 and Table S3), a feature of scRNA-seq measuring thousands of transcripts but not spatial proteomics measuring tens of proteins.

To assess the generalizability of our comparative analyses, we extended these analysis to CODEX measurements of 24 proteins in 220,155 cells from 30 islets in a non-diabetic donor (Tables S9 and S10). Similar to IMC results (Figures 2 and S2), qualitative and quantitative studies showed higher performance of AnnoSpat in predicting endocrine cell types with high quality antibodies from CODEX data compared to other algorithms (Figure S3 and Table S11, and Supplemental Notes for AnnoSpat Benchmarking). Together, these comparative analyses indicated the advantage of using AnnoSpat for accurate, comprehensive, and rapid cell-type annotation from IMC and CODEX spatial proteomic measurements.

### AnnoSpat improves accuracy of cell type identification in expert-annotated pan-creata

To further demonstrate AnnoSpat’s ability in accurate cell-type annotation, we compared AnnoSpat and expert-annotated endocrine cell composition in pancreata of non-diabetic and diabetic donors^16^ (Figure 3 and Table S12). Comparison of AnnoSpat- and expert-annotated cells revealed concordance in endocrine cell composition in 12 out of 15 (80%) examined IMC samples (Figures 3A and 3B). Notably, our analysis revealed manual cell-type mislabeling in the remaining three discordant samples (Figures 3C, 3D and S4). Compared to expert annotation, AnnoSpat identified markedly higher and lower percentages of PP and delta cells in (HPAP002, Head) and (HPAP015, Head), respectively (Figures 3A and 3B). Close examination of these samples confirmed the accuracy of AnnoSpat cell-type annotation and showed high expression of canonical PP cell marker protein, PPY, in AnnoSpat-annotated cells (Figures 3C, 3D, S4A, and S4B). While AnnoSpat identified a high percentage of alpha cells in the body region of HPAP006 pancreas, manual annotation indicated a low percentage of alpha and a high percentage of delta cells (Figures 3A and 3B). In line with AnnoSpat cell-type annotation, we observed a higher percentage of cells with elevated levels of glucagon (a canonical marker of alpha cells) in HPAP006 pancreas body (Figures S4C and S4D).

**Figure 3:**
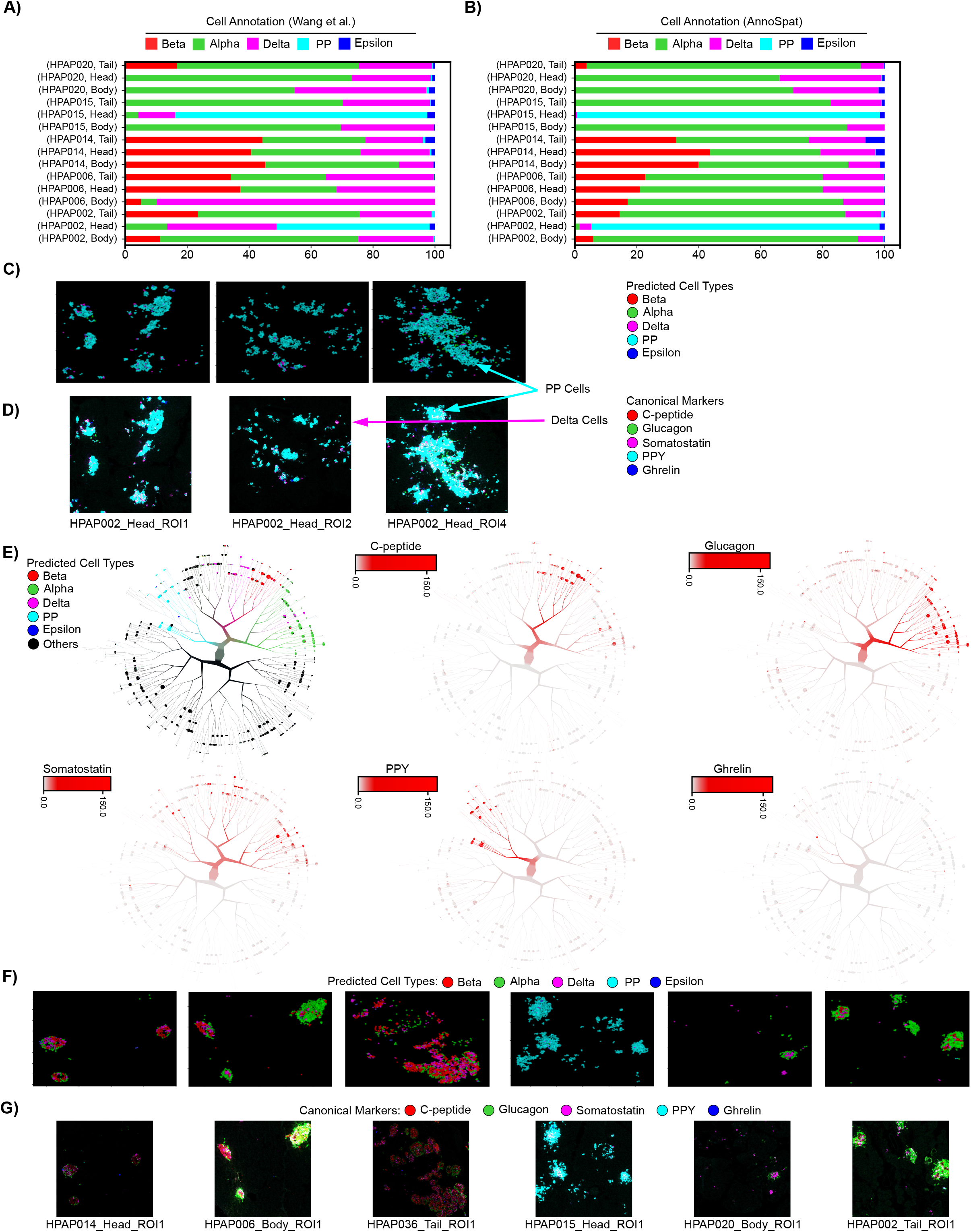
Comparison of expert and AnnoSpat endocrine cell-type annotation. **(A, B)** Proportion of expert-annotated (A) and AnnoSpat-annotated (B) endocrine cell types from the IMC of different pancreas regions of donors studied in^16^. **(C, D)** Representative protein channel intensities (expression levels) from IMC images of donors with discordant expert and AnnoSpat cell-type annotation is overlaid with AnnoSpat predicted cell types (C) or endocrine canonical marker protein channels (D). **(E)** From left to right, top to bottom: TooManyCells tree overlaid by AnnoSpat-predicted cell types, and expression levels of c-peptide, glucagon, somatostatin, pancreatic polypeptide protein (PPY), and ghrelin marking beta, alpha, delta, PP, and epsilon cells, respectively in *n* = 65, 643 cells across *m* = 141 slides of 16 pancreas donors. **(F, G)** Six representative IMC images from *m* = 141 slides of 16 donors overlaid by AnnoSpat-predicted cell types (F) or endocrine canonical marker protein channels (G).

Given the single-cell resolution of IMC data, we next used various visualization methods to compare the AnnoSpat-assigned cell types with canonical marker protein expression levels in individual endocrine cells. Uniform manifold approximation and projection (UMAP) plots of AnnoSpat cell label and endocrine marker protein expression clearly visualized specificity of glucagon, c-peptide, somatostatin, ghrelin, and PPY expression in cells labeled as alpha, beta, delta, epsilon, and PP cells, respectively (Figure S5). A similar analysis using TooManyCells, which visualizes cell-cell protein expression relationships as a tree^13^, further confirmed our UMAP analysis and demonstrated a high association between AnnoSpat-predicted endocrine cell types and expression of their canonical marker proteins at cell clusters (Figure 3E).

Finally, we used the locational information from the spatial proteomic data to directly compare AnnoSpat annotations and marker protein intensities of endocrine cells *in situ*. This analysis revealed a stark concordance between the position of cells predicted as alpha, beta, delta, epsilon, and PP and the intensity of glucagon, c-peptide, somatostatin, ghrelin, and PPY expression on randomly selected IMC and CODEX slides, respectively (Figures 3F, 3G, S3I, and S3J). This single-cell resolution analysis complemented benchmarking against expert-annotated samples and further demonstrated the accuracy of AnnoSpat in identifying the identity of individual cells in spatial proteomic data.

### AnnoSpat showed PP cell count increase during T1D progression

Linking expression of canonical protein markers with the predicated cell types demonstrated AnnoSpat’s superior ability to automatically identify various cell types within the heterogeneous pancreas tissue, the site of T1D pathogenesis (Figures 2 and 3). To further evaluate AnnoSpat functionality, we next examined whether it could correctly detect progressive changes in pancreata during T1D progression. We thus compared IMC data of four non-diabetic (control) and four diabetic (T1D) donors with data of eight donors with autoantibodies towards islet proteins (AAbs) but without T1D medical history (AAb^+^) (Table S2). Given that many T1D patients harbor AAbs in their bloodstream prior to clinical diagnosis, we postulated that this analysis might elucidate pathogenic events prior to disease manifestation.

Control, AAb^+^ and T1D donors demonstrated distinct total normalized protein expression patterns in cell types annotated by AnnoSpat (Figure 4A). Comparison of cell-type composition revealed marked decreases in beta-cell counts of T1D donors (Figure 4B), as expected^17,18^. This analysis further showed a notable increase in the number of cells labeled as PP in T1D donors (Figure 4B).

**Figure 4:**
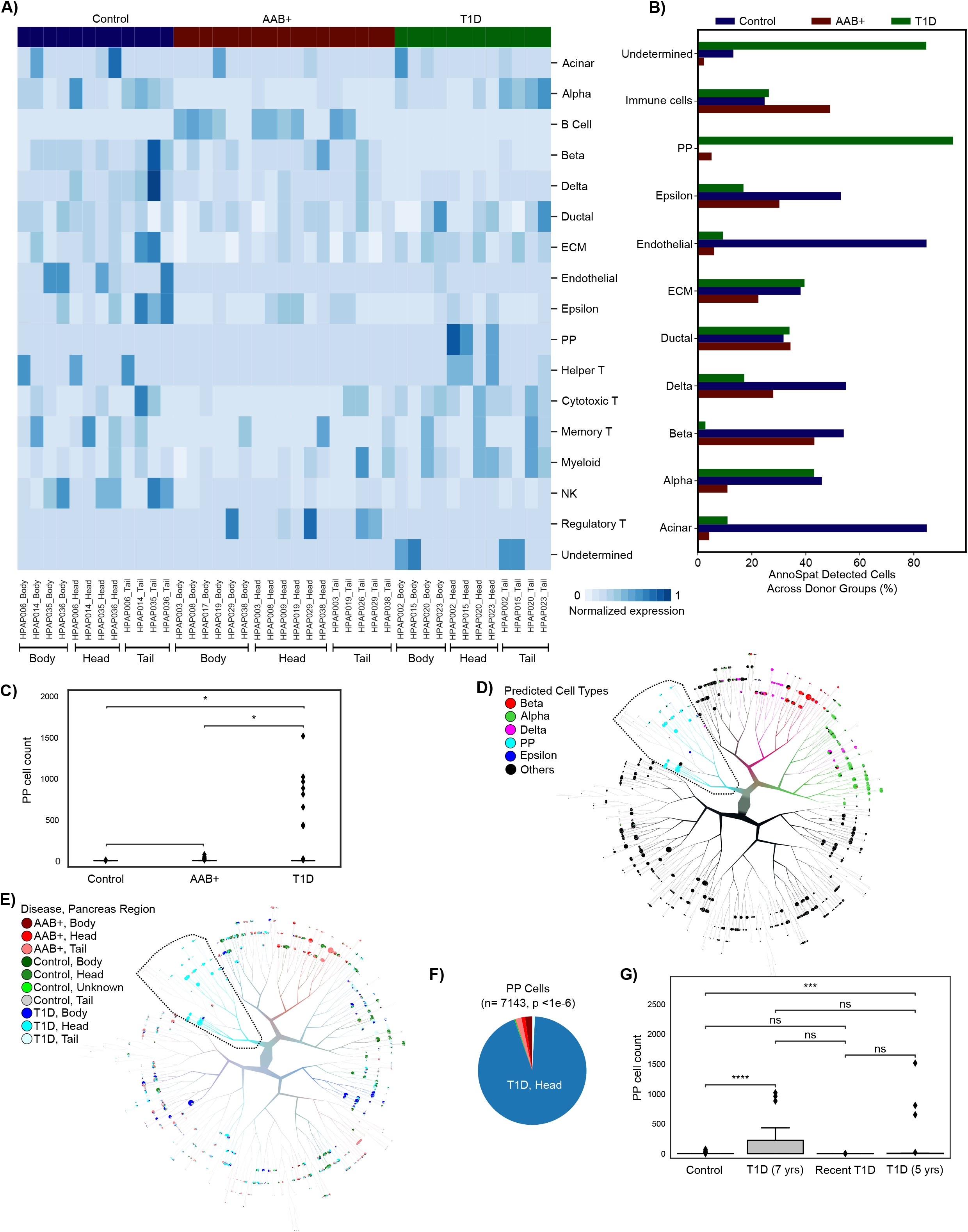
PP cell count increases in the pancreas head during T1D progression. **(A)** Heatmap showing total normalized protein expression for each pancreas region across non-diabetic control, T1D and AAb^+^ donors. Normalized protein expression for each cell type is calculated by scaling for ROI count per donor pancreas region (3/*ROIcount*) of min-max and TF-IDF normalized expression levels. **(B)** Bar plots showing percentage of each AnnoSpat-annotated cell type across pancreata of control, T1D, and AAb^+^ donors. **(C)** Plots showing PP cell counts in pancreata from control, T1D, and AAb^+^ donors. **(D, E)** TooManyCells tree overlaid with AnnoSpat-predicted cell types (D), as well as disease status and pancreas region (E). TooManyCells default parameters (quartile normalization and filter threshold of channel intensity < 250 and marker protein intensity <1) were used. **(F)** Pie chart showing fraction of PP cells from different pancreas regions across control, T1D, and AAb^+^ cohorts. **(G)** Box-and-whisker plots quantifying PP cell counts in control and T1D donors stratified by disease duration. ** *p*-value <0.01, *** *p*-value<0.001, **** *p*-value<0.0001, n.s. not significant (*p*-value ≥ 0.05). Box-and-whisker plots: center line, median; box limits, upper (75^th^) and lower (25^th^) percentiles; whiskers, 1.5 · interquartile range; points, outliers.

In contrast to beta cells, the role of PP cells in T1D etiology is less understood. Furthermore, there are conflicting reports regarding changes in the PP cell count during T1D development^19–24^. We thus compared the number of PP cells identified within the pancreata from control, AAb^+^, and T1D donors. This analysis showed a marked increase in the number of PP cells in T1D pancreata (Figure 4C), as reported^16,21^.

To further scrutinize this observation, we examined the location of individual AnnoSpat-annotated endocrine cells (Figure 4D) on the TooManyCells tree of non-diabetic control and T1D pancreatic cells (Figure 4E). This single-cell resolution analysis further showed that AnnoSpat-annotated PP cells were disproportionately located at T1D pancreas heads (Figures 4E and 4F), with the exception of HPAP020. Given AAb^+^ donors also did not show elevated PP-cell counts (Figure 4C), we tested whether disease progression correlates with changes in PP-cell numbers. PP cell counts were comparable in control and T1D donors with less than 5 years of T1D, and were markedly lower than donors with a prolonged T1D (Figures 4G and S6). Notably, fewer PP cells were found in the head of HPAP020 pancreas, a 14-year-old donor who, with missed T1D diagnosis, passed away within days of T1D onset (Figures 4G and S6). To further substantiate this observation, we closely examined data from Damond et al.^25^. This data set confirmed our observation and showed enrichment of PP cells in the only donor with long-duration of T1D and available head section sample in this cohort (nPOD case 6,264). Together, these data showed the ability of AnnoSpat to identify rare PP cells, and further suggest changes in the PP cell count during T1D progression in our cohort, which could be absolute or relative, respectively, due to PP cell hyperplasia or PP-cell poor region atrophy impacting tissue sampling.

In addition to tissue level analysis (Figure 4), IMC data can be used for single-cell resolution study of protein expression changes in T1D. To this end, we sought to identify the proliferating cell populations within pancreatic tissue using Ki67 as a protein marker. Average normalized protein levels showed high Ki67 expression in various immune populations (Figure 5A and Materials and Methods). To identify the proliferating cell types and their disease status, we used the TooManyCell tree to identify individual Ki67^+^ cells (Figure 5B). This analysis revealed that myeloid and regulatory T cells comprised most of the Ki67^+^ cells (Figure 5C). Examination of highly proliferating cells’ positions further revealed that these cells were disproportionately located in the tail region of AAb^+^ and T1D pancreata (Figure 5D). Although the role of these highly proliferating immune cells in T1D patients awaits further investigation, this analysis demonstrated the ability of AnnoSpat to simultaneously stratify multiple cell types enabling detailed molecular phenotyping to identify changes in the immune milieu of complex diseases such as T1D.

**Figure 5:**
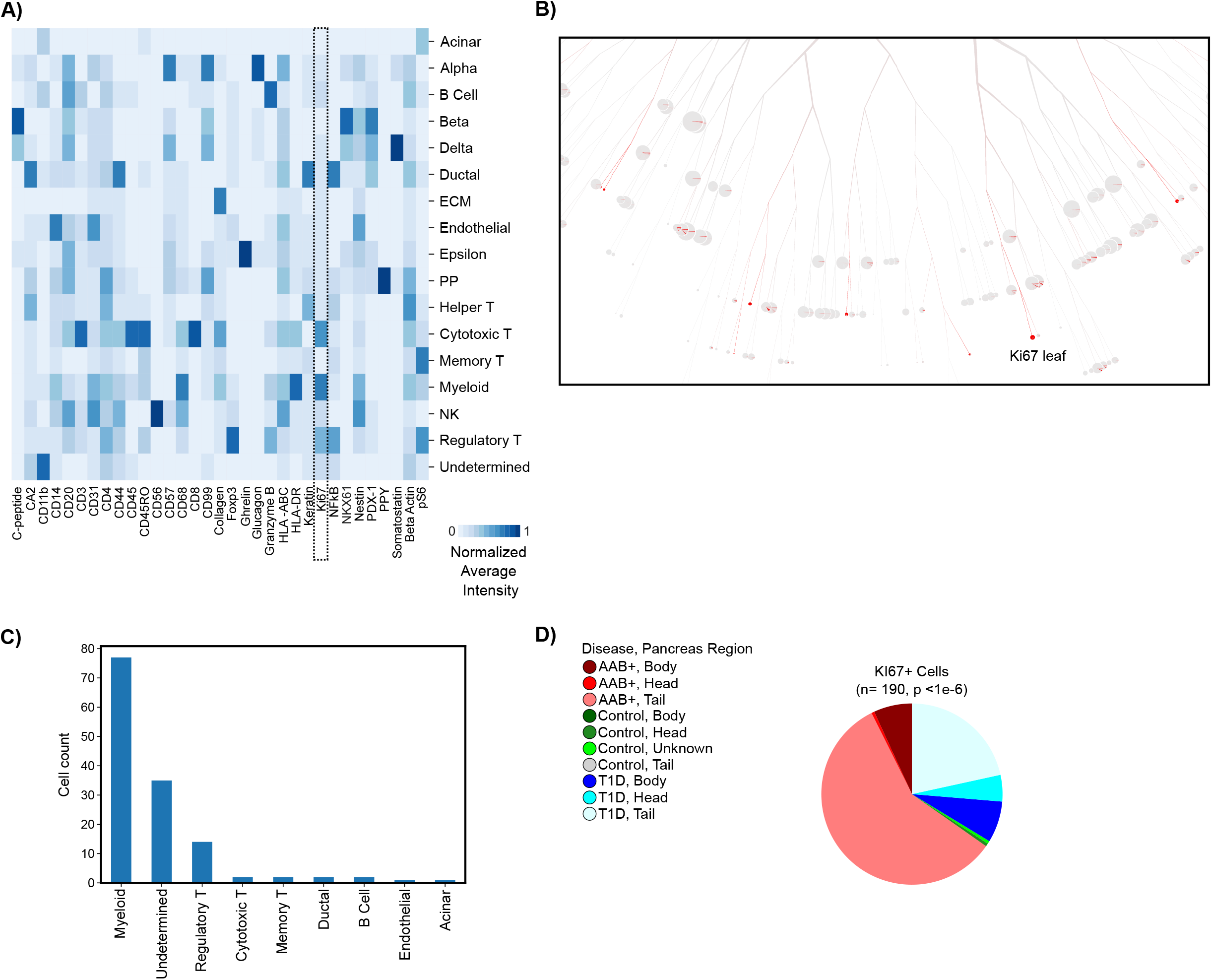
Myeloid and regulatory T cells are hyper-proliferative in T1D pancreata. **(A)** Heatmap showing normalized average expression of 33 IMC-measured proteins across AnnoSpat-annotated cell types. Dash-lined box marking Ki67 column. **(B)** TooManyCells sub-tree colored by Ki67 expression. **(C)** Bar plots showing cell-type count of *n* = 190 cells within the TooManyCells sub-tree in (B). **(D)** Pie chart showing fraction of Ki67^+^ cells from different regions of pancreata of control, T1D, and AAb^+^ donors (*p*-value: chi-square test).

### AnnoSpat elucidates CD8^+^ T cell infiltration in islet during T1D development

Having identified composition of endocrine cells in control, AAb^+^, and T1D samples, we next sought to understand the spatial relationships between islets and immune cells (Figure 6). To quantify cell proximity, we used AnnoSpat’s ‘Spatial Pattern Finder’ functionality, which identifies spatial patterns of cells by reporting cross-correlation functions from point process theory. Briefly, AnnoSpat interprets each cell as a point in space with the cell type as a discrete feature “mark”. In this space, AnnoSpat measures the expected number of cells per unit area. AnnoSpat compares this number, which is its null model, to the expected number of cells for a given cell-type pairing to find whether these cell types tended to aggregate across a range of distances (Figure 1C and Materials and Methods). To compare mark cross-correlation functions between distributions of ROIs, we proposed multiple measures to summarize mark cross-correlation functions into single values such as the distance at the maximum correlation value for each ROI.

**Figure 6:**
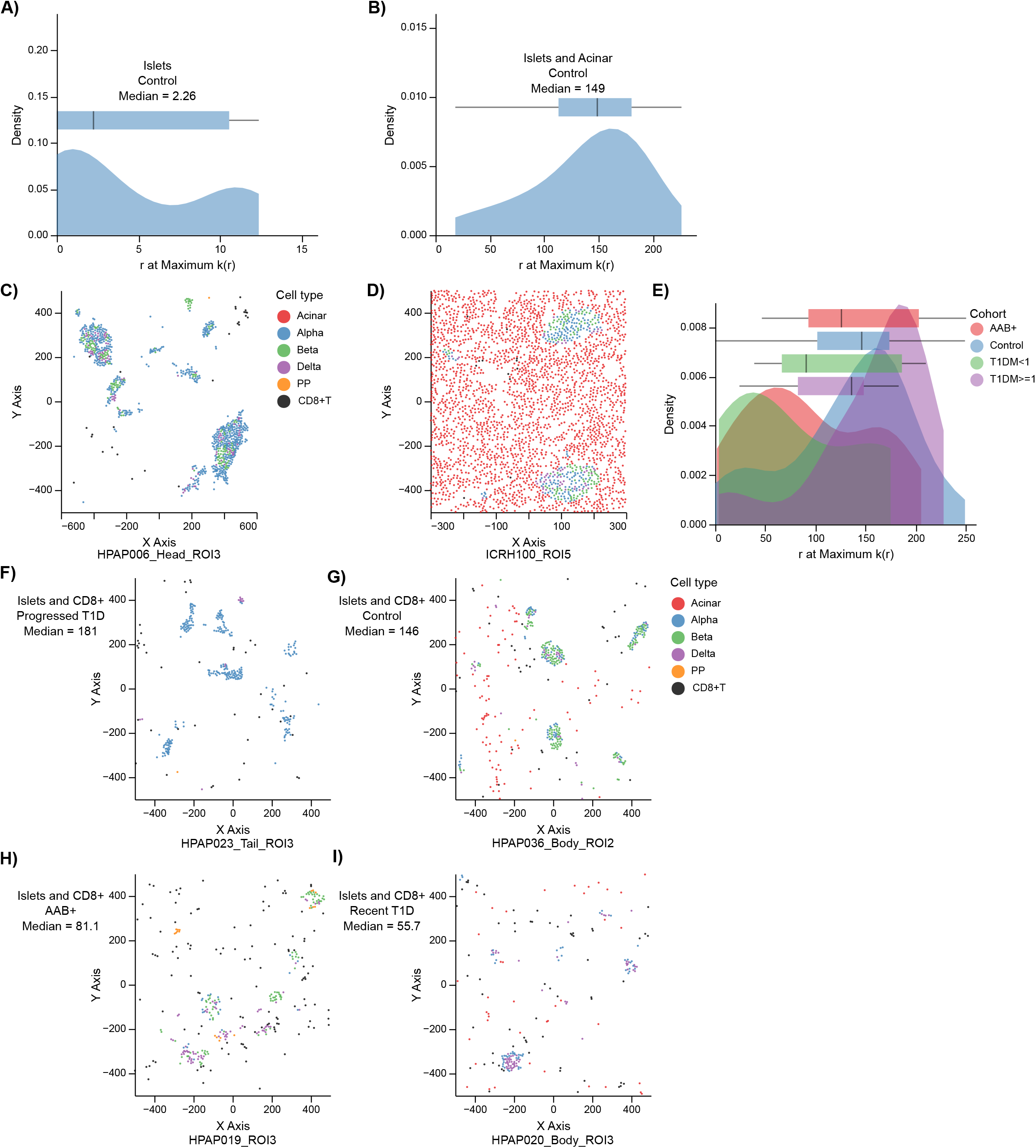
The extent of CD8^+^ T cell infiltration in islets changes during T1D progression. **(A, B)** Distributions with box-and-whisker plot overlays of distance *r* at the maximum value of *k*(*r*) across all ROIs for endocrine cells with respect to themselves (A) or with respect to acinar cells (B). **(C, D)** Scatter plots showing location of cells within ROIs at the median of (A) and (B) distributions are plotted at (C) and (D), respectively. Cells are colored by AnnoSpat-predicted cell types. Endocrine cells tend to aggregate around themselves more often than with acinar cells. **(E)** The distributions with box-and-whisker plot overlays of the distance at the maximal point in the mark cross-correlation functions across control, AAb^+^, recent T1D, and prolonged T1D. AAb^+^ and recent T1D tend to have greater aggregation of islets with CD8^+^ T cells than control and prolonged T1D cohorts. **(F-I)** Scatter plots showing location of cells within ROIs at the median of each cohort in (E). From lowest to highest aggregation: prolonged T1D (F), control (G), AAb^+^ (H), and recent T1D (I). Cells are colored by AnnoSpat-predicted cell types. Box-and-whisker plots: center line, median; box limits, upper (75^th^) and lower (25^th^) percentiles; whiskers, 1.5 · interquartile range; points, outliers.

To verify the use of mark cross-correlation functions in IMC data, we first used AnnoSpat’s Spatial Pattern Finder to compare endocrine cell aggregation into islets with their aggregation with acinar cells in the ROIs of the control donors. As expected, endocrine cells aggregated more with each other (Figure 6A, median 2.26 distance at maximum correlation value) than with acinar cells (Figure 6B, median 149). These spatial relationships were confirmed by visual inspection of the samples present at the median values, where the endocrine cells were generally aggregated with each other while positioned more randomly with respect to acinar cells (Figures 6C and 6D). Using an alternative measure to summarize the mark cross-correlation functions, we observed similar spatial patterns confirming the use of both measures in comparison of cell-cell proximity patterns (Figures S7A-D and Materials and Methods).

To further examine the utility of AnnoSpat’s Spatial Pattern Finder in studying T1D pathogenesis, we next quantified the spatial relationship between CD8^+^ T and islet cells. Given that the destruction of insulin-producing beta cells by cytotoxic CD8^+^ T cells contributes to T1D pathogenesis^17,18^, we tested the hypothesis that T1D progression would be characterized by different levels of cytotoxic CD8^+^ T cell infiltration in islets. Applying mark cross-correlation functions to all ROIs for four cohorts of control, AAb^+^, recent T1D (< 1 year), and prolonged T1D (≥1 year) revealed two distinct patterns of spatial relationships between islets and CD8^+^ T cells: AAb^+^ with recent T1D and control with prolonged T1D (Figures 6E-I). Non-diabetic control donors, as expected, had relatively low degrees of CD8^+^ T cell infiltration in islets (median 146). Similarly, we observed low levels of CD8^+^ T infiltration in islets of prolonged T1D (median 181). In contrast, both AAb^+^ (median 81.1) and recent T1D (median 55.7) had markedly higher aggregation of CD8^+^ T cells within islets relative to both control and prolonged T1D groups (Kruskal-Wallis *p*< 0.01), suggesting potential differences in immune responses during T1D progression (Figures 6E-I). Furthermore, AAb^+^ and recent T1D tissues showed similar levels of CD8^+^ T cells infiltration in islets (*p*> 0.05), suggesting similar autoimmune responses in early stages of T1D with and without clinical diagnosis (Figures 6E-I). These differential spatial relationships were confirmed using our alternative mark cross-correlation summarization measure (Figures S7E-I). Visual inspection of IMC images further supported these quantitative observations (Figure S7J-M), suggesting that CD8^+^ T cell infiltration in islets increases in early onset but not prolonged T1D.

## Discussion

Spatial profiling of cells *in situ* has enabled comprehensive exploration of cellular organization in tissues. Such high-throughput data has led to the need for automated cell-type annotation and methods to quantify cell-cell spatial relationships. However, current methods for cell-type annotation in spatial proteomic analysis either involve manual labeling, which prohibits scalability, or suffer from low accuracy as shown in our comparative studies. To address this unmet need and overcome these limitations, we developed AnnoSpat for efficient and accurate prediction of individual cell types and their relationships within spatial proteomic data. Using both quantitative and qualitative benchmarking, we demonstrated that AnnoSpat can rapidly and accurately predict the identity of millions of cells in complex human pancreata profiled with IMC and CODEX assays. Our comparative studies further showed that AnnoSpat can predict lineages of large fraction of cells with high accuracy, while other existing cell annotation algorithms failed to do so. AnnoSpat accuracy is further exemplified by identifying endocrine cell populations undetected by manual annotation.

Using the unique capabilities of AnnoSpat, we accurately recapitulated known changes in the pancreas microenvironment during T1D progression such as depletion of beta cells with minimal manual intervention on a dataset of over a million cells. Moreover, our analysis supported the possibility of changes in the number of PP cells within the pancreas head region during T1D progression. We also observed proliferating immune cells were enriched within the tail region of pancreata from AAb^+^ and T1D donors. Differential immune-cell heterogeneity during T1D progression was not solely limited to cell count. By using AnnoSpat’s spatial relationship quantification functionality, we found different spatial patterns between immune and endocrine cells across donor types. Specifically, AnnoSpat reported marked changes in CD8^+^ T cell infiltration in islet during T1D progression, suggesting alternative disease categorizations – donors recently diagnosed along with AAb^+^ donors versus control donors and those with prolonged T1D, potentially due to a reduced autoimmunity response after beta cell depletion in prolonged T1D.

AnnoSpat is generalized for spatial signle-cell proteomics, potentially applicable to many tissue types and disease conditions. Yet, the performance of AnnoSpat and other for automated cell type annotation algorithms could be impacted by IMC and CODEX antibody quality, such as the ones used for PPY and CD4 in CODEX and IMC experiments here. To enhance usability across domains, AnnoSpat is well documented and available as an easy-to-install standalone program through pip at https://github.com/faryabiLab/AnnoSpat. We also provided Annospat’s spatial pattern quantification functionality as part of the TooManyCells suite located at https://github.com/GregorySchwartz/too-many-cells.

## Materials and Methods

### Supplemental Notes for AnnoSpat Benchmarking

Our comparative analysis presented in Figures 2 and S2 highlighted the intricacy of differences in the ability of AnnoSpat, semi-supervised clustering (SSC), SCINA, AUCell, and Astir to identify endocrine cell types in pacreatic tissues. Here, we presented a more comprehensive description of these differences. Figures 2A, 2B, and 2C row 1; as well as Tables S4, S5, and S6 revealed the algorithms’ cell-type- and disease-status-related performance differences. Despite a comparable performance in detecting most cells, AUCell exhibited low accuracy in identifying beta cells in T1D samples, where immunological destruction of beta cells results in low beta-cell abundance. Most methods underperformed in detecting the epsilon cells, which is a rare endocrine cell type in islets. SCINA and AUCell more accurately detected PP compared to delta cells. AnnoSpat and SSC more accurately detected delta cells in the control samples compared to other algorithms. SCINA, designed for scRNA-seq count data, underperformed on both metrics and sample conditions, underscoring the need for cell-type calling algorithms specifically designed for spatial proteomics data that is fundamentally different from scRNA-seq count data. Importantly, Astir, developed for cell-type detection from IMC data, showed lower accuracy in identifying many cell types in both control and T1D samples.

Inspection of protein expression profiles of cells annotated as alpha, beta, PP, delta, and epsilon from IMC of T1D and non-diabetic donors (Figures 2A, 2B, and 2C, rows 2 to 6) further confirmed our quantitative benchmarking and showed the higher performance of AnnoSpat compared to SCINA and Astir, the other method specifically designed for cell-type annotation from spatial proteomic data. AnnoSpat- and Astir-predicted beta cells from T1D samples, where beta cells are rare, showed high levels of immune cell-restricted proteins CD57 and HLA-ABC. Comparing the result of cell-type prediction from T1D alone with T1D plus control cohorts (i.e. Combined) showed that additional samples improved the performance of AnnoSpat more-so than Astir. Notably, Astir equally failed to detect epsilon cells in T1D, control, and combined data sets. CD11b, a marker of dendritic cells, was the highest expressed protein in the Astir-predicted delta cells. Furthermore, Astir-predicted alpha cells expressed high levels of somatostatin, a canonical marker of delta cells. Similar to Astir, SCINA failed to detect delta cells in samples from non-diabetic donors. Moreover, SCINA-annotated PP cells were less homogeneous compared to the AnnoSpat-labeled cells.

In addition to endocrine cells, AnnoSpat effectively detected other cell types that had high quality antibodies and are commonly present in the pancreatic tissue (Figures S1C and S2). For instance, AnnoSpat clearly identified CD8^+^ T cells that had a specific antibody (Figures S1C and S2). Conversely, detection of helper and memory T cells was less accurate due to their less specific antibodies (Figures S1C and S2).

We further extended our comparative studies to CODEX measurements of 24 proteins in 220,155 cells from 30 islets in a non-diabetic donor (Tables S9 and S10). For this analysis, we focused on detection of alpha, beta, and delta cells due to lower quality of PPY and grehlin antibodies (Figure S3A). SI and DB metrics suggested that AnnoSpat’s performance was comparable to AUCell and SCINA for most populations (Figure S3B, Table S11). Yet, a close examination of labeled cells revealed that in contrast to AnnoSpat, SCINA-annotated beta cells expressed high levels of somatostatin, a canonical marker of delta cells (Figures S3C and S3F). While AnnoSpat identified a pure delta cell population, SCINA-annotated delta cells lacked high levels of canonical marker SST. AUCell-annotated delta cells expressed high levels of CD206, ARG1, and CD4, canonical markers of macrophages and T helper cells, respectively (Figures S3C and S3G). In contrast to IMC analysis, AnnoSpat consistently outperformed SSC in predicting abundant endocrine cells from CODEX data. For instance, ghrelin, a canonical marker of epsilon cells, was highly expressed in SSC-labeled delta cells (Figure S3C and S3D), supporting advantage of ELM usage in AnnoSpat. Similar to benchmarking with IMC data (Figure 2), AnnoSpat outperformed Astir in predicting endocrine cells from CODEX data (Figures S3C and S3E). Besides beta cells, Astir failed to annotate other major endocrine cell populations (Figure S3E). Close examination of data further showed high levels of non-beta-cell-associated proteins in Astir-labeled beta cells (Figures S3B and S3E). Notably, we observed high levels of canonical marker proteins in the nucleus and/or cytoplasm of AnnoSpat-labeled cells from high-resolution CODEX data, further supporting the accuracy of AnnoSpat in cell type annotation (Figure S3H). Together, these comprehensive analyses indicated the advantage of using AnnoSpat for accurate, comprehensive, and rapid cell-type annotation from IMC and CODEX spatial proteomic measurements.

### IMC and CODEX Data

IMC data were obtained from Formalin-Fixed Paraffin-Embedded (FFPE) pancreatic tissues collected by the Human Pancreas Analysis Program (HPAP) as described previously^16^, and is available at the HPAP data repository https://hpap.pmacs.upenn.edu/. CODEX data were obtained from the same source, and will be deposited at the HPAP data repository. In IMC, cell segmentation of all images was performed with the Vis software package (Visiopharm). All image channels were pre-processed with a 3 × 3-pixel median filter. Afterwards, cells were segmented by applying a polynomial local linear parameter-based blob filter to the Iridium-193 DNA channel of each image to select objects representing individual nuclei. Identified nuclear objects were restricted to those greater than 10 *μm*^26^. The detected objects were dilated up to seven pixels to approximate cell boundaries. For all proteins, the average pixel intensity of the channel per cell was exported from Visiopharm and used for AnnoSpat’s input. Cell locations on each ROI were also exported for AnnoSpat’s input.

### AnnoSpat Overview

AnnoSpat is a tool to annotate single cells from their proteomic profiles and measure spatial cellular relationships using their *in situ* coordinates within the ROI. AnnoSpat takes as input a single-cell raw proteomic data with associated spatial information as well as a signature file listing potentially both positive and negative protein signatures associated with desired cell types. The format of the signature file can be found in Tables S3 and S10.

AnnoSpat first normalizes the protein channel intensity data to reduce the effect of outliers and varied protein intensity scales (Materials and Methods: Data Processing). AnnoSpat then randomly splits the normalized data into two partitions (training and testing sets). Cells from 50% of each ROI are placed in the training set, while the remaining are used as the testing set. If the ROIs’ disease condition/status is available, AnnoSpat can split the ROIs by status to ensure that equal percentage of each type of ROIs are included in each set.

AnnoSpat implements constrained K-means semi-supervised clustering^27,28^ to identify groups of cells in the training set that are similar in proteomic space. AnnoSpat’s constrained K-means clustering is initialized by “initial cluster centroids”, providing “cell-type aware” clustering (Materials and Methods: Generation of Initial Cluster Centroids). The number of clusters is deterministic and is equal to *K* + 1, where *K* denotes the number of expected cell types in the sample. The additional (*K* + 1)th cluster accounts for other cell types in the experiment that are not specified in the marker file, including “Unknown” ones. The output of constrained K-means produces the cells that are predicted to be related and thus are used by AnnoSpat as a training set to learn the label of the remaining cells by training an extreme learning machine classifier (ELM)^12^ (Materials and Methods: Training Extreme Learning Machine Classifier). The trained model is saved to label cells from other data sources, eliminating the need for re-clustering or re-training whenever new data is available.

AnnoSpat can use the cell-type labels and cellular coordinates to quantify spatial relationships between each pair of cell types (Materials and Methods: AnnoSpat’s Spatial Pattern Finder). Briefly, AnnoSpats uses point process theory to quantify relationships (aggregation or repulsion) between any two cell types across a range of distances. This information is summarized with a variety of different metrics including the distance at the maximum correlation, the distance at which the correlation first becomes positive or negative, and more in order to quantify proximity relationships across ROIs. Interactive plots of each cell location with observed feature (protein expression) distributions are also outputted to facilitate data exploration (For example see Figures 6 and S7).

### AnnoSpat Data Processing

To reduce the effect of outliers, AnnoSpat first calculates Data matrix *D* by log transforming cell-by-protein channel intensity (expression) after addition of pseudo-count 1. Specifically, *d_c×p_* = *e_c×p_* + 1, where *e_c×p_* is the expression of protein *p* channel in cell *c*. Then, AnnoSpat unit normalizes the log-transformed intensity matrix to scale each cell vector to unit length. This projects each cell to a unit sphere in the proteomic space. We denote the normalized proteomic matrix by *X* obtained from scaling each row *d_i_* of *D* as follows:

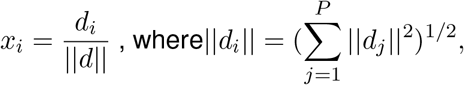

where,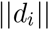 denotes the *l*_2_ or Euclidean norm of *i* th cell. *P* is the number of measured proteins.

This step accounts for variable expression across proteins and correlates the Euclidean distances (used for clustering) between cell vectors and cosine distances in the proteomic space. Compared to euclidean distance, the angle between the cell vectors in proteomic space better reflects cell-cell similarities/differences^29^.

### Generation of Initial Cluster Centroids

As opposed to traditional K-means where the initial cluster centroids are randomly selected, AnnoSpat implements constrained K-means that follows a more “cell-type aware” approach^27,28^. Initial cluster centroids are obtained from representatives of each cluster (“cell-type” here). AnnoSpat calculates initial cluster centroids by taking the mean of representative cells *R_k_* for each cluster *k* = 1*,…, K* + 1. The number of clusters is one more than the number of cell types *K*; an extra (“Unknown”) cluster accounts for cell types not included in the marker file.

AnnoSpat obtains the cluster representations *R*_1_, *R*_2_*,…, R_K_*_+1_ by:

1. Obtaining positive and negative markers *M* ^+^ and *M^−^* from the signature file.
2. Calculating the score *M_c_* for *c^th^* cell type by multiplying the protein intensities corresponding to positive markers and the compliment of protein intensities corresponding to negative markers as follows:

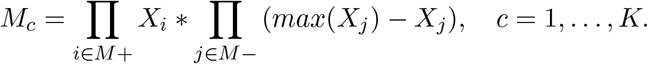
3. Selecting cell representatives *R*_1_, *R*_2_*,…, R_K_* of cell types *c* = 1*,…, K* in the signature file such that they have

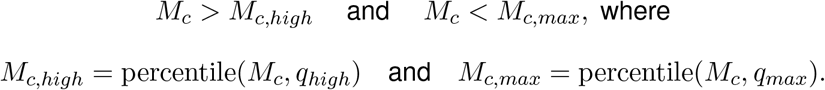 The value *q_high_* is adaptive and can be optionally chosen based on prior knowledge of the number of cells from the cell type present in the data (defaulting to the 95*^th^* percentile). Here, *q_high_* was set to 99 ≤ *q_high_* ≤ 99.9 and 99.5 ≤ *q_high_* ≤ 99.99 for various cell types in the analysis of pancreas IMC and CODEX data, respectively. *M_c,high_* is the score cut-off to pick cluster representative cells as the ones having a very high score *M_c_* corresponding to the *c^th^* cell type. The threshold *q_max_* is set to 100 or a value slightly less than that to make sure that assay artifacts are not included in the initial cluster centroid calculation. Here, *q_max_* was set to 99.999 and 100 for the analysis of pancreas IMC and CODEX data, respectively.
4. Assigning cell representatives *R_K_*_+1_ of the “Unknown” cluster such that they have *M_c_* < *M_c,low_* where

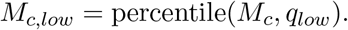 The threshold *q_low_* defines the cut-offs to pick cells with expression <*M_c,low_* in all cell types.

AnnoSpat performs the above procedure to assign cluster representative cells in decreasing order of cell-type abundance (representative of more abundant cell types are selected first). Cell-type abundance acts as a proxy for the expected number of cells for each cell type and is obtained by summing cell intensities of the scale-normalized canonical protein markers.

Once the cell representatives *R*_1_, *R*_2_*,…, R_K_*_+1_ have been assigned, AnnoSpat computes initial centroids 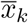 for cluster *k* = 1*,…, K* + 1 by taking the average across the representative cells *R_k_* as follows:

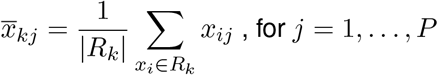

where *x_ij_* represents the intensity of the *j^th^* protein in *i^th^* cell.

### Cell Labeling with Semi-supervised Clustering

AnnoSpat takes the cell representatives *R_k_*’s and initial cluster centroids 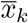’s and iteratively runs constrained K-means algorithm on the cells from 50% of the ROIs included in the training set as shown in Algorithm 1. *L_i_* denotes the cluster label assigned to the *i^th^* cell and *C_k_* denotes the set of cells in cluster *k*. The assigned cell labels are the predicted cell types of training data for the AnnoSpat’s Annotator.

### Training Extreme Learning Machine Classifier

AnnoSpat uses the cell-type labels *L* predicted by its semi-supervised clustering algorithm as training labels *Y_T R_* to then learn an ELM classifier^12^. The classifier predicts the label of remaining cells in new ROIs not included in the training data. We implemented ELM in AnnoSpat because it is a single-layer feed-forward neural network classifier and does not need to be iteratively tuned via backpropagation. This would enable AnnoSpat to learn accurate cell type prediction models markedly faster than gradient-based learning techniques. AnnoSpat’s ELM is implemented as follows:

**Algorithm 1** Constrained K-means

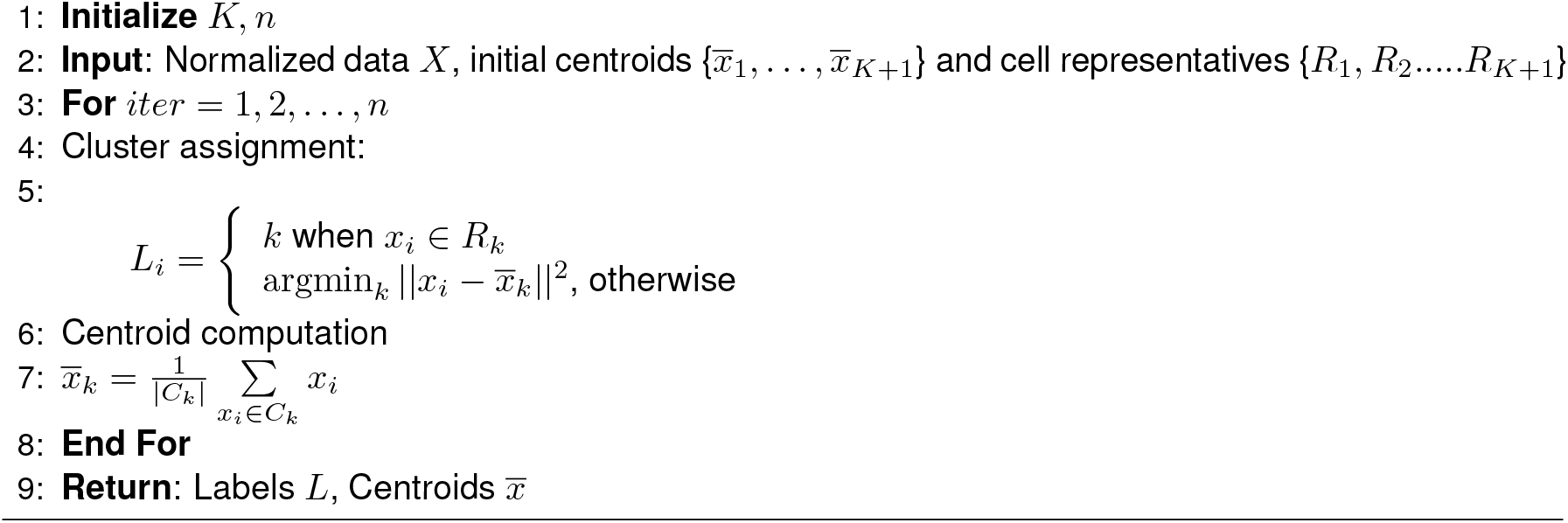

- Assign input layer weights *W_I_* and bias *b_I_* randomly from normal distributions:

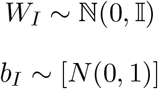
- Compute hidden layer output *H*:

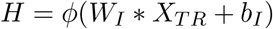 Here, *ϕ* denotes the activation function used at the hidden layer, and *X_T R_* is the normalized protein intensity of training set.
- Compute the output layer weights *W_O_*

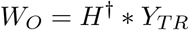

Here *H*^†^ is the Moore–Penrose inverse of hidden layer output matrix H. The training labels *Y_T R_* are transformed into a one-hot encoded format to avoid ordinal relationship interpretability between cell types by the model.

Once the output weights are learned, the types (labels) of new cells *Y_T S_* can be predicted from their normalized protein expression *X_T S_* by the learned weights in ELM:

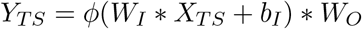

### Data Processing for Visualization

Data for Heatmaps in Figures 2 and S1C have been normalized to penalize the expression of non-specific proteins using an analog variant of TF-IDF (term frequency-inverse document frequency) normalization after min-max scaling of protein expression. The specificity of a protein can be quantified as an inverse function of the number of cell types in which it is expressed (its abundance across various cell types). Hence, the normalized value of protein *p_i_* is obtained by multiplying each value by the logarithm of ratio of total protein abundance *p_total_* in the data and the abundance of that protein across all cell types *p_sum_*. If *p* is the expression of protein *p_i_* in cell *c_j_*, then the normalized value is calculated by:

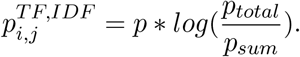

In min-max normalization, min and max values are the 0.01^*th*^ and 99.99^*th*^ percentile expression, respectively.

### AnnoSpat’s Spatial Pattern Finder: Quantification of Cell Proximity Pattern

In order to quantify the relationships between cell types in the T1D pancreas, we interpreted the cell locations and cell type labels as a marked point pattern. A point pattern provides the locations of observations; here, cell locations are represented as Cartesian coordinates. Each cell can have additional features known as marks; here, each cell’s mark is the predicted cell type. By realizing the marked point pattern as a random marked point process, we can quantify cell type spatial relationships. A point process is a random set of points, where the number of points and their locations are both random. Using point process theory, we can understand the relationship between cell types not as a single index, but rather as many values resulting in formulation of a given function of distance *r*.

The standard model of a point process ≩ assumes that the process extends all space, but the observed region is bounded by a window *W*. Then we can define the data as an unordered set^30^

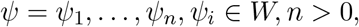

the point pattern of **Ψ**.

Now we can define our ROI within the context of marks. Consider the marked point pattern as an unordered set of cells observed within a window *W* with marks in *M*

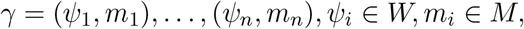

where *ψ_i_* is the location and *m_i_* is the mark of cell *i*, respectively^30^. Marks may be continuous real numbers, such as cell size, or discrete, such as cell type. Our objective is to quantify the dependence between the marks of two cells of distance *r* apart in the marked point process **Γ**. This dependence, known as the mark correlation function *k_f_* (*r*), is informally defined as^30,31^

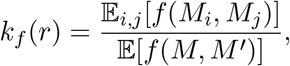

where *M_i_, M_j_* are marks of two cells separated by distance *r*, *M, M′* are independent realizations of the marginal distribution of marks, and E is the *intensity* of a point process, or the average density of points (the expected number of points per unit area), and where E_*i,j*_ is the conditional expectation that there exist cells at locations *i* and *j* separated by distance *r*. While *f* is any function that returns a non-negative real value, we commonly use *f* (*m*_1_, *m*_2_) = *m*_1_*m*_2_ for continuous marks and 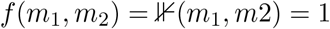 where *m*_1_ = *m*_2_ and = 0 for everything else for discrete (categorical) marks^30^. Then, *k_f_* (*r*) = 1 suggests a lack of correlation such that under random mark labeling, *k_f_* (*r*) ☰ 1. The interpretation of greater than or less than 1 would be determined by the chosen function *f*, but throughout this study we interpret > 1 as correlated and < 1 as anti-correlated. This mark correlation function, however, assumes that cell type would be a single mark and does not specify the relationship between, for instance, CD8^+^ T cells and islet cells.

To understand the relationship between any two cell types, we expand the mark correlation function *k_f_* (*r*) to define the mark cross-correlation function, *k_mm_*(*r*). Here, instead of *m_i_* ∈ *M* as a single mark, we define ⋗_*ia*_ ∈ *M* as the value of mark *a* in cell *i* from the row vector of marks ⋗_*i*_ attached to cell *i*. Instead of a single mark for cell type, we convert the mark into a mark row vector *m_i_* for cell *i* containing *c* entries, where each index 0 <*j* ≤ *c* represents an indicator value for cell type *a*. In short, ⋗_*ia*_ = 1 indicates that the cell *i* is of cell type *a*.

Using this expanded mark vector, we can define the mark cross-correlation function^30^ as

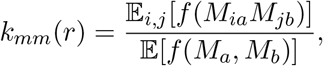

where *M_ia_* and *M_jb_* are the marks *a* and *b* attached to cells *i* and *j*, respectively, while *M_a_* and *M_b_* are independent random values drawn from all cells at mark indices *a* and *b*, respectively. Here, *f* is defined as with the mark correlation function. Using categorical marks for cell types, we then interpret *k_mm_*(*r*) > 1 as correlated, < 1 as anti-correlated, and = 1 as random. We carried out all mark cross-correlation analyses using the spatstat R package^30^.

The output of each mark cross-correlation function on an ROI is a series of correlation values as a function of distance *r*. To compare across several ROIs, we summarized each curve by either the *r* at the maximum *k_mm_*(*r*) (max*_r_ k_mm_*(*r*)) (Figure 6) or the log-transformed ratio of the maximum *k_mm_*(*r*) to the *r* at the maximum *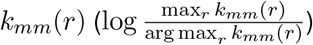* (Figure S7). The former value decreases with increasing aggregation (the highest correlation is with cells with smaller *r*) while the latter increases with increasing aggregation. To compare distributions, we used Kruskal-Wallis one-way analysis of variance for multiple hypotheses followed by pairwise Mann-Whitney *U* tests.

## Acknowledgments

This work was supported in part by R01-CA230800, R01-CA248041 (to R.B.F.), Canada Research Chairs Program (to G.W.S.), Human Islet Research Network (RRID:SCR-014393), and Human Pancreas Analysis Program (RRID:SCR-016202) through DK112217, DK123594, DK104211, DK108120, and DK112232.

## Authors Contributions

Conceptualization: R.B.F., G.W.S.; Methodology: A.M., G.W.S., R.B.F.; Software: A.M., G.W.S.; Investigation: A.M., G.W.S., R.B.F.; Formal Analysis: A.M.,D.T., D.S., G.W.S., R.B.F.; Resources and Reagents: R.B.F., Y.J.W., A.C.P, K.H.K., G.V., A.N.; Writing-Review & Editing: G.W.S., R.B.F.; Writing-Original Draft: A.M., G.W.S., R.B.F.; Supervision: R.B.F.; Funding Acquisition: R.B.F., G.V., A.N.

## Data Availability

IMC and CODEX data are available at PANC-DB, the data portal of Human Pancreas Analysis Program (HAPA) developed by the Faryabi Lab, https://hpap.pmacs.upenn.edu/.

## Code Availability

AnnoSpat is available at https://github.com/faryabiLab/AnnoSpat. Spatial Pattern Finder is also available as part of the TooManyCells suite located at https://github.com/faryabib/too-many-cells.

## Supplementary Materials

### Supplementary Figures

**Figure S1:**
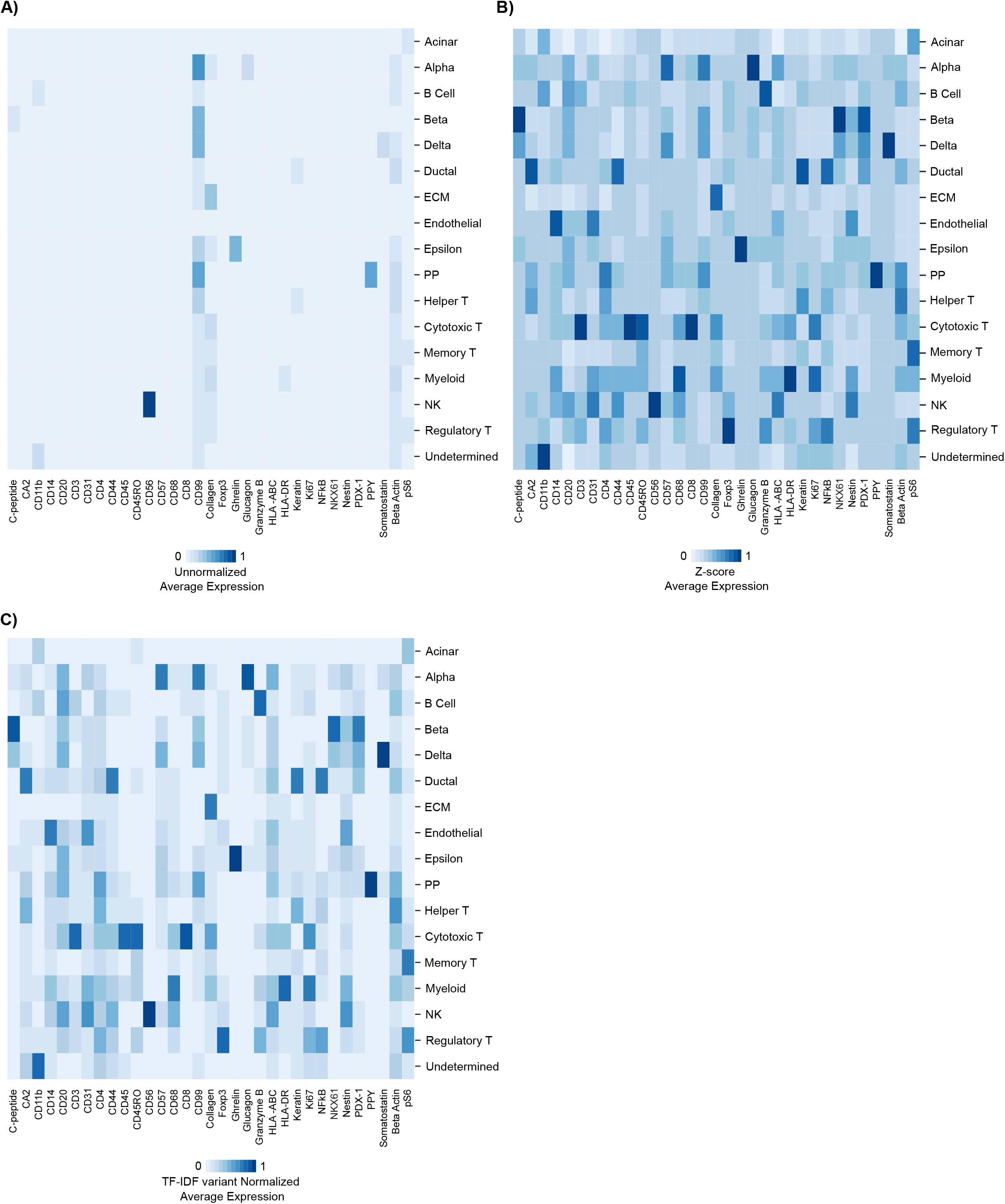
Comparison of various IMC data normalization methods. Heatmaps showing average unnormalized **(A)**, protein-wise z-score normalized **(B)**, and a variant of TF-IDF normalized **(C)** expression levels of 33 IMC-measured proteins across AnnoSpat-annotated cell types. Heatmaps comparison indicates the benefit of a variant of TF-IDF for normalization in visualizing continuous protein expression readouts. Note: TF-IDF variant normalization is only used for data visualization and not cell-type annotation.

**Figure S2:**
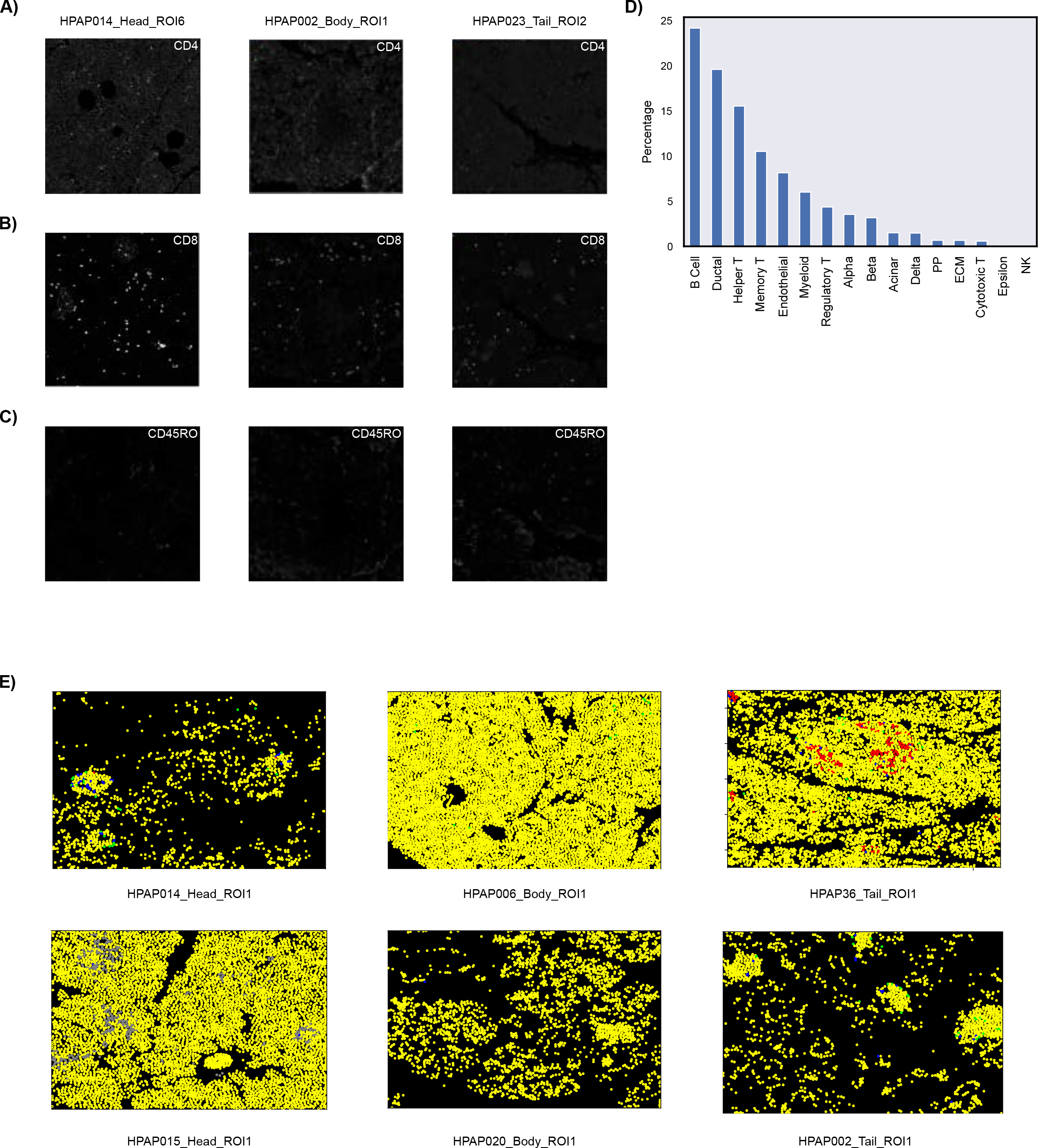
Comparison of AnnoSpat and AUCell cell-type annotation. **(A-C)** Randomly selected IMC images of ROIs from pancreas head, body, tail comparing CD4 (A), CD8 (B) and CD45RO (C) staining quality showing higher quality of CD8 compared to CD4 and CD45RO staining. **(D)** Bar plots showing percentage of AnnoSpat-annotated cell types that AUCell failed to annotate. **(E)** Yellow pseudo-color marking AnnoSpat-annotated cells that AUCell failed to annotate on randomly selected IMC images. Other cell types are colored as before (e.g. refer to Figure 3C).

**Figure S3:**
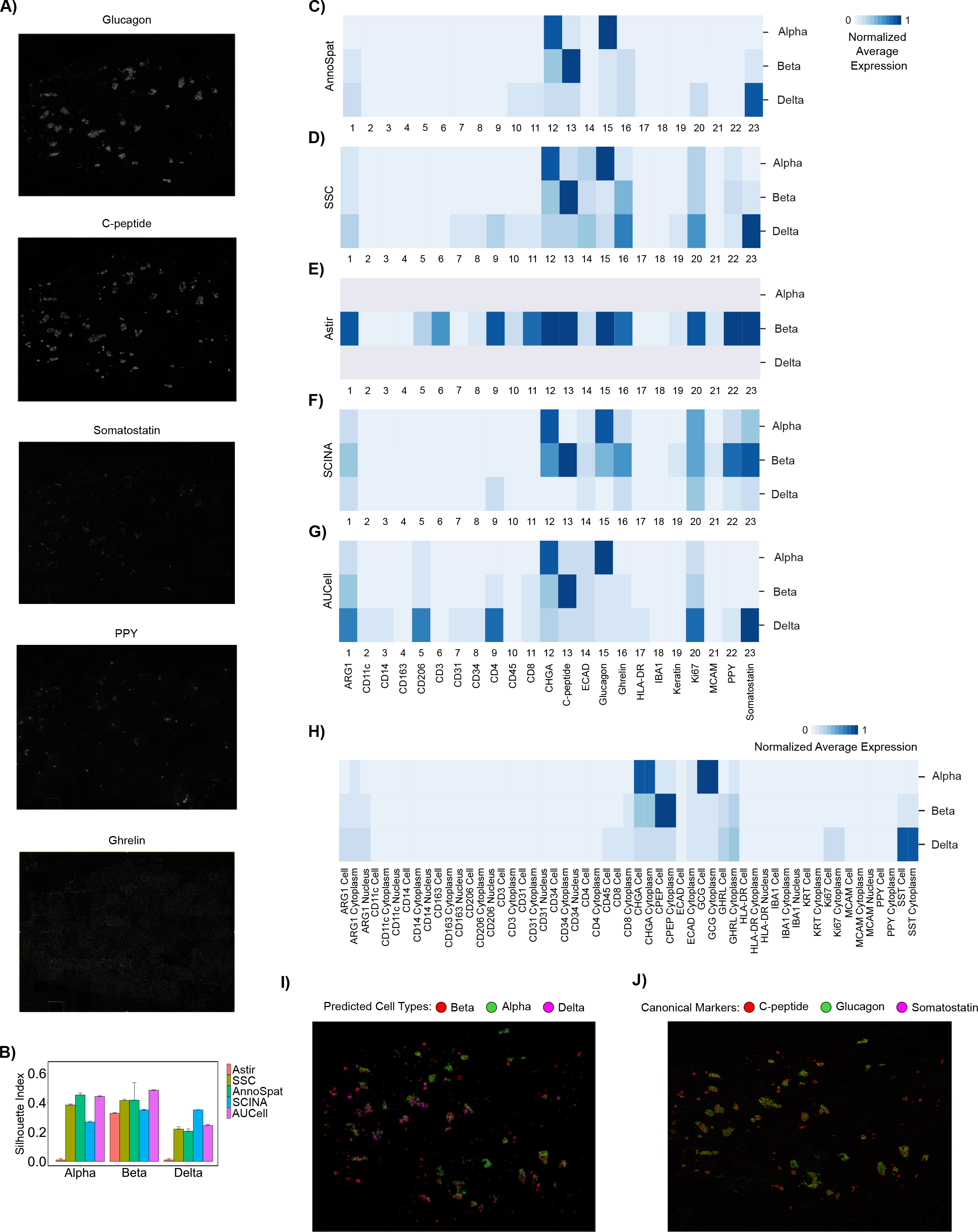
Comparative analysis of AnnoSpat cell-type annotation from CODEX data. **(A)** From top to bottom: raw images of glucagon, c-peptide, somatostatin, PPY, and ghrelin showing non-specificity of PP and epsilon markers in CODEX experiments. **(B)** Bar plots with error bars showing average and standard deviation Silhouette Index (SI) for cells annotated as alpha, beta, and delta by AnnoSpat, semi-supervised clustering (SSC), Astir, SCINA, and AUCell from non-diabetic pancreas CODEX data (*m* = 10 sets of *n* = 50, 000 cells randomly selected from *n* = 220, 155 measured cells). **(C-G)** Heatmaps showing marker proteins’ normalized average expression levels for cells labeled as alpha, beta, and delta by AnnoSpat, SSC, Astir, SCINA, and AUCell from non-diabetic pancreas CODEX data (*n* = 220, 155 measured cells). **(H)** Heatmap showing marker proteins’ normalized average expression levels separately for the nucleus and cytoplasm of the cells annotated as alpha, beta, and delta by AnnoSpat based on protein intensity in the entire cells from non-diabetic pancreas CODEX data (*n* = 220, 155 measured cells). **(I, J)** CODEX image is overlaid by AnnoSpat predicted cell types (I) or alpha (glucagon), beta (c-peptide), and delta (somatostatin) marker protein channels (J).

**Figure S4:**
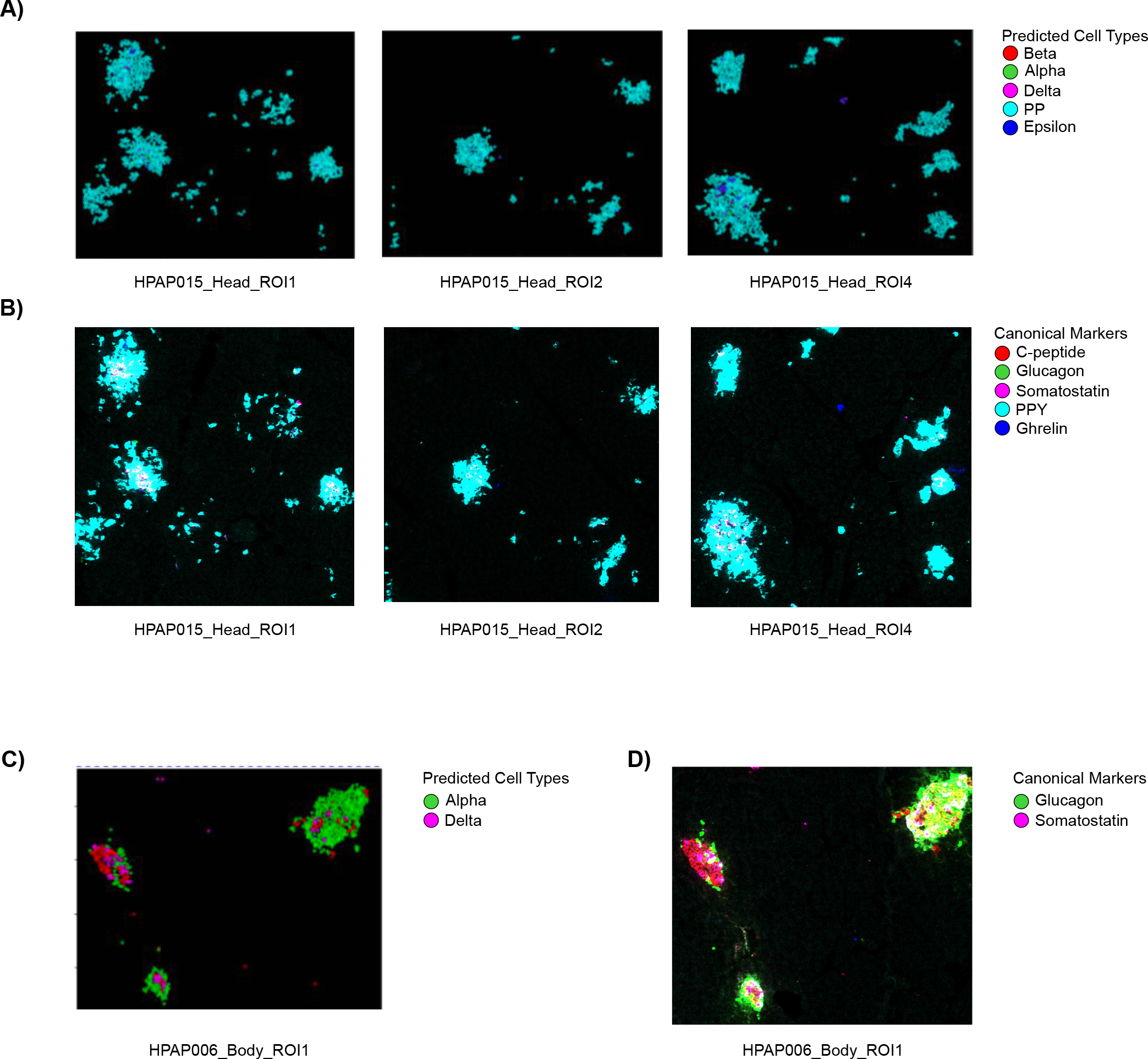
Comparison of AnnoSpat and expert annotation of pancreatic endocrine cell types. **(A-D)** Representative IMC images from donors with discordant expert and AnnoSpat cell-type annotation in Figures 3A and 3B are overlaid with AnnoSpat-predicted cell types (A and C) or endocrine canonical marker protein channels (B and D). C-peptide, glucagon, somatostatin, PPY, and ghrelin marking beta, alpha, delta, PP, and epsilon cells, respectively.

**Figure S5:**
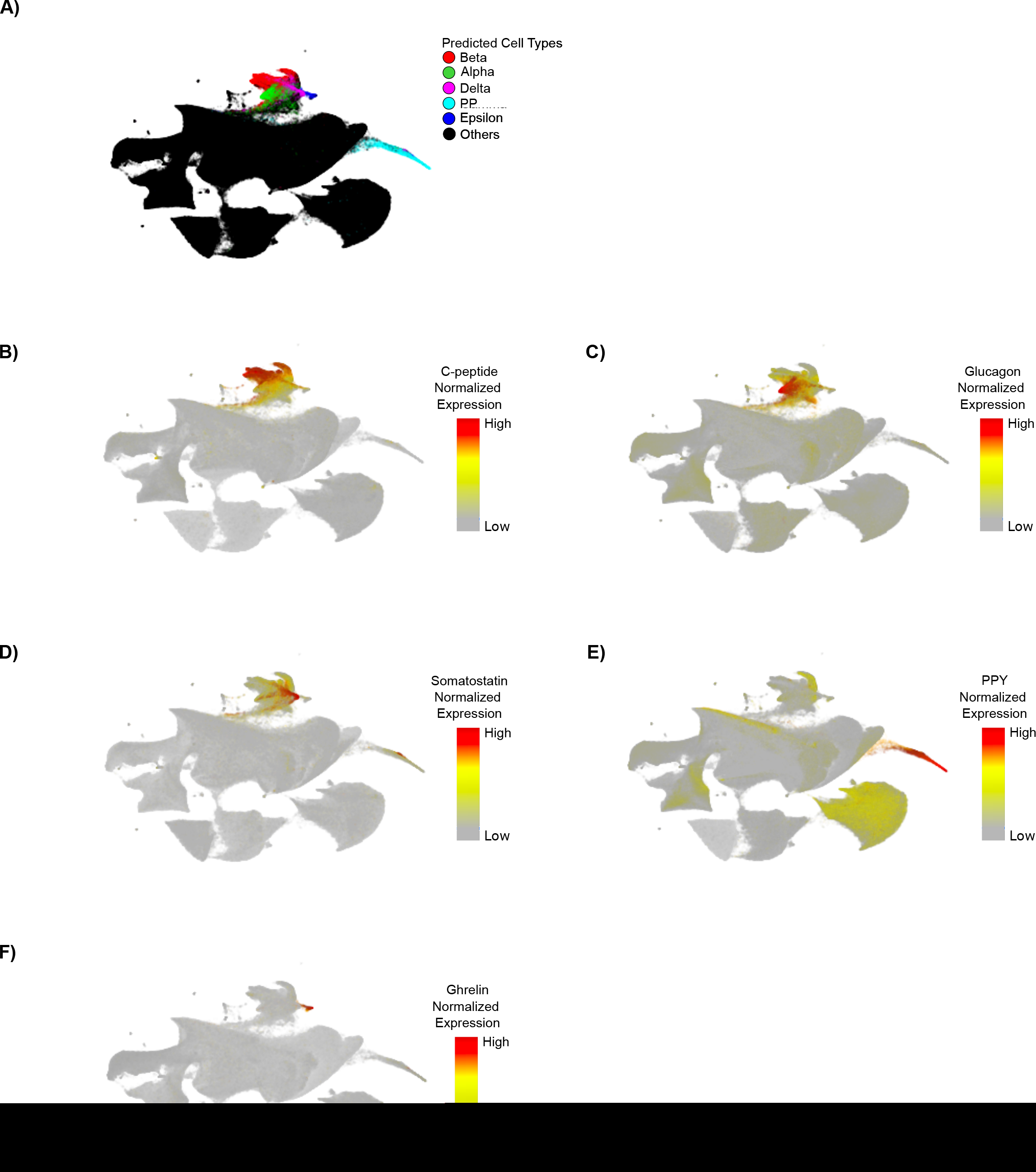
Comparison of protein marker expression levels and AnnoSpat annotations across pancreatic endocrine cell types. **(A-F)** UMAP plots overlaid by AnnoSpat-predicted cell types (A), and expression levels of c-peptide (B), glucagon (C), somatostatin (D), PPY (E), and ghrelin (F) in *n* = 65, 643 cells across *m* = 141 slides of 16 pancreas donors.

**Figure S6:**
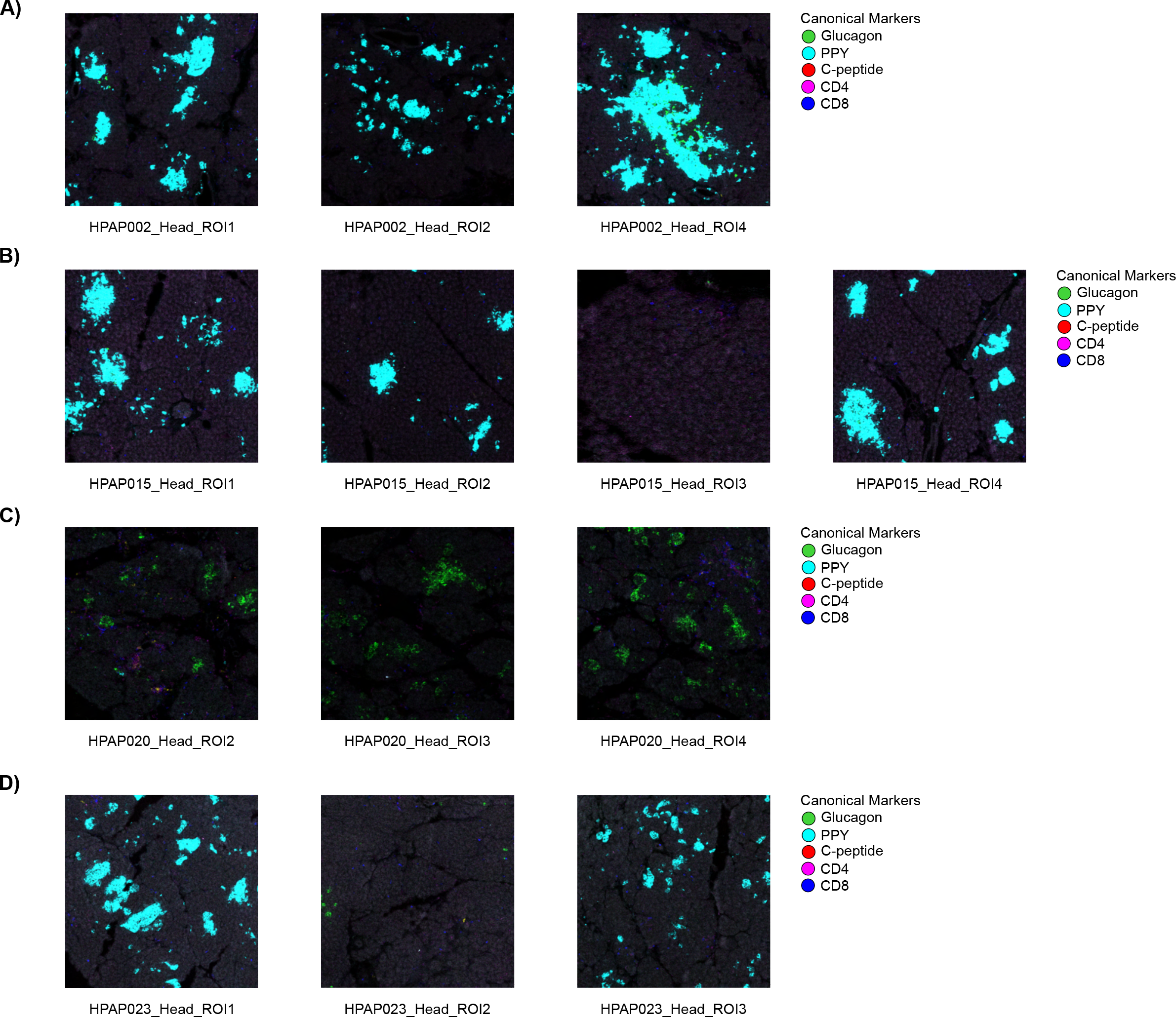
PP cell count increases in the pancreas head during T1D progression. **(A-D)** IMC images from pancreatic head ROIs overlaid with expression levels of canonical protein markers of alpha (glucagon), beta (c-peptide), PP (PPY), helper T (CD4), and cytotoxic T (CD8) cells.

**Figure S7:**
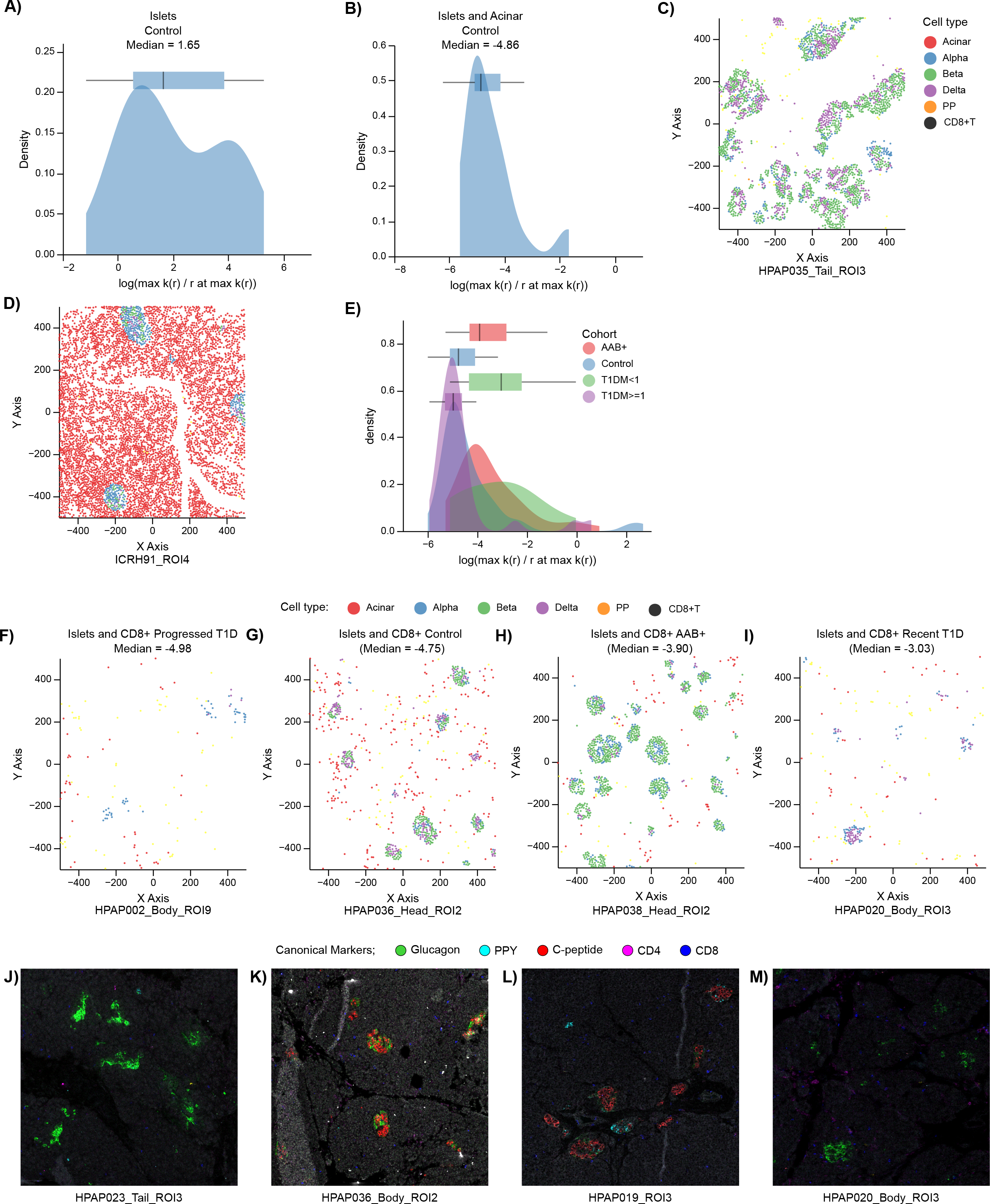
The extent of CD8^+^ T cell infiltration in islets changes during T1D progression. Analysis corresponding to Figure 6 but with an alternative summarization measure of mark cross-correlation function, which takes into account correlation value as well as distance (*r*):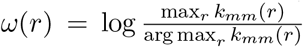 **(A, B)** Distributions with box-and-whisker plot overlays of *ω*(*r*) *r* across all ROIs for endocrine cells with respect to themselves (A) or with respect to acinar cells (B). **(C, D)** Scatter plots showing location of cells within ROIs at the median of (A) and (B) distributions are plotted in (C) and (D), respectively. Cells are colored by AnnoSpat-predicted cell types. Endocrine cells tend to aggregate around themselves more often than with acinar cells. **(E)** The distributions with box-and-whisker plot overlays of *ω*(*r*) across control, AAb^+^, recent T1D, and prolonged T1D. AAb^+^ and recent T1D tend to have greater aggregation of islets with CD8^+^ T cells than control and prolonged T1D cohorts. **(F-I)** Scatter plots showing location of cells within ROIs at the median of each cohort in (E). From lowest to highest aggregation: prolonged T1D (F), control (G), AAb^+^ (H), and recent T1D (I). Cells are colored by AnnoSpat-predicted cell types. **(J-M)** IMC images from pancreatic ROIs overlaid with expression levels of canonical protein markers of alpha (glucagon), beta (c-peptide), PP (PPY), helper T (CD4), and cytotoxic T (CD8) cells confirming changes in the CD8^+^ T cell infiltration in islets during T1D progression. Images in (J) to (M) correspond to scatter plots in Figures 6F to 6I, respectively. Box-and-whisker plots: center line, median; box limits, upper (75^th^) and lower (25^th^) percentiles; whiskers, 1.5 · interquartile range; points, outliers.

## Supplementary Tables

**Table S1**

IMC antibody panel.

**Table S2**

HPAP pancreas donor information.

**Table S3**

AnnoSpat’s marker file input for annotating the listed cell types from IMC antibodies.

**Table S4**

SI and DB scores for labeling endocrine cells from IMC samples of T1D donors’ pancreata. Numbers in parentheses: standard deviation. NA: no cell was annotated.

**Table S5**

SI and DB scores for labeling endocrine cells from IMC samples of non-diabetic (control) donors’ pancreata. Numbers in parentheses: standard deviation. NA: no cell was annotated.

**Table S6**

SI and DB scores for labeling endocrine cells from IMC samples of combined non-diabetic and T1D (Combined) donors’ pancreata. Numbers in parentheses: standard deviation. NA: no cell was annotated.

**Table S7**

Fraction of endocrine cells labeled by each algorithm from IMC samples of T1D, non-diabetic (control), and combined T1D and control (Combined) donors’ pancreata.

**Table S8**

Mean and standard deviation of run-time for listed algorithms to annotate cells from IMC samples of T1D, non-diabetic (control), and combined T1D and control (Combined) donors’ pancreata. Each algorithm was run three times on data sets of *n* = 374, 397, *n* = 795, 604, and *n* = 1, 170, 001 cells from IMC samples of T1D, control, and combined T1D and control donors using a machine with Ubuntu 20.04, 1.05TB Memory, Intel Xeon Gold CPU 6230R @ 2.1GHz, 2 physical processors 52 cores, and 104 threads.

**Table S9**

CODEX antibody panel.

**Table S10**

AnnoSpat’s marker file input for annotating the listed cell types from CODEX antibodies.

**Table S11**

SI and DB scores for labeling alpha, beta, and delta cells from a non-diabetic donor pancreas CODEX. Numbers in parentheses: standard deviation. NA: no cell was annotated.

**Table S12**

Fraction of expert-annotated endocrine cell types in different regions of pancreata from donors studied in^16^.

